# Simultaneously monitoring aquatic and riparian biodiversity using riverine water eDNA

**DOI:** 10.1101/2020.06.20.162388

**Authors:** Haile Yang, Hao Du, Hongfang Qi, Luxian Yu, Xindong Hou, Hui Zhang, Junyi Li, Jinming Wu, Chengyou Wang, Qiong Zhou, Qiwei Wei

## Abstract

Environmental DNA (eDNA) metabarcoding for biodiversity monitoring is a critical technical advance. Both aquatic and terrestrial biodiversity information can be detected in riverine water eDNA. However, it remains unverified whether riverine water eDNA can be used to simultaneously monitor aquatic and terrestrial biodiversity. Our specific objective was to assess the effectiveness of monitoring aquatic and riparian biodiversity using riverine water eDNA. We proposed that the monitoring effectiveness (the proportion of aquatic and terrestrial biodiversity information detected by riverine water eDNA samples) could be approximated by the transportation effectiveness of land-to-river and upstream-to-downstream biodiversity information flow. We conducted a case study in a watershed on the Qinghai-Tibet Plateau and estimated the effectiveness of using riverine water eDNA to monitor aquatic and riparian biodiversity based on comparing the operational taxonomic units (OTUs) and species assemblages of three taxonomic communities detected in riverine water eDNA samples and riparian soil eDNA samples in spring, summer, and autumn. The aquatic and riparian biodiversity of a watershed on the Qinghai-Tibet Plateau could be simultaneously effectively monitored using riverine water eDNA on summer or autumn rainy days. Monitoring bacterial communities was more efficient than monitoring eukaryotic communities. On summer rainy days, 43%-76% of riparian species could be detected in water eDNA samples, 92%-99% of upstream species could be detected in a 1-km downstream eDNA sample, and more than 50% of dead bioinformation (i.e., the bioinformation labeling the biological material without life activity and fertility) could be monitored 4-6 km downstream for eukaryotes and 13-19 km for bacteria. We encourage more studies on the monitoring effectiveness for each taxonomic community in other watersheds with different environmental conditions. We believe that in future ecological research, conservation and management, we could efficiently monitor and assess the aquatic and terrestrial biodiversity by simply using riverine water eDNA samples.

## 1 Introduction

Biodiversity monitoring is the basis of ecological research, biodiversity conservation and ecosystem management (Cardinale et al., 2012; Hooper et al., 2012; Dixon et al. 2019; Pawlowski et al. 2020). Traditional biodiversity monitoring methods are cost- and time-consuming and require high levels of expertise, in which biodiversity is often studied from a local and low spatio-temporal resolution perspective and is generally not available at a wide taxonomic breadth, high spatio-temporal resolution and large spatio-temporal scale (Anderson, 2018; Altermatt et al., 2020; Pawlowski et al. 2020). This limits the development of ecological research, biodiversity conservation and ecosystem management. Currently, metabarcoding and high-throughput sequencing of environmental DNA (eDNA, DNA extracted from environmental samples such as water, soil, and air) provide novel opportunities to monitor biodiversity (Rodgers and Mock 2015; Deiner et al., 2016; Ushio et al. 2017; Carraro et al., 2018; Gogarten et al. 2019; Johnson et al. 2019; Seeber et al. 2019; Pawlowski et al. 2020; Sales et al. 2020). As an efficient and easy-to-standardize non-invasive monitoring approach (Thomsen & Willerslev, 2015; Deiner et al. 2016; Lugg et al., 2018; Ravindran,2019; Seymour, 2019; Shogren et al. 2019), and with the continuous advancements in DNA sequencing technology, using eDNA metabarcoding to monitor biodiversity is an appropriate method to revolutionize biodiversity monitoring by enabling the census of wide taxonomic species on a high spatio-temporal resolution and large spatio-temporal scale (Thomsen & Willerslev, 2015; Bass et al. 2015; Deiner et al., 2016; Valentini et al., 2016; Bálint et al. 2018; Cristescu & Hebert, 2018; Altermatt et al., 2020).

Streams and rivers connect upstream and downstream regions, connect land with waterbodies, and transport materials and information through extensive and heterogeneous network systems (Luo et al., 2011; Deiner et al., 2016; Wang et al., 2016; Shogren et al., 2017; Matsuoka et al., 2019). Riverine water eDNA incorporates biodiversity information across terrestrial and aquatic biomes, and offers a novel and spatially integrated component that can be used to simultaneously monitor aquatic and terrestrial biodiversity (Deiner et al., 2016; Matsuoka et al., 2019). Therefore, a sample of riverine water eDNA has the potential to simultaneously monitor both aquatic and terrestrial biodiversity of a watershed for biodiversity research, conservation, and management. However, its viability and monitoring effectiveness (the proportion of aquatic and terrestrial biodiversity information that could be detected using limited riverine water eDNA samples) has not been systematically verified.

The biodiversity monitoring effectiveness of riverine water eDNA depends on the land-to-river and upstream-to-downstream transportation effectiveness of the bioinformation (eDNA) labeling terrestrial and upstream biodiversity (Deiner & Altermatt, 2014; Deiner et al., 2016; Jerde et al. 2016; Sansom & Sassoubre, 2017; Pont et al., 2018). To verify the approach of using riverine water eDNA to simultaneously monitor aquatic and terrestrial biodiversity, we could estimate its monitoring effectiveness accordingly to assess the land-to-river and upstream-to-downstream transportation effectiveness of the corresponding bioinformation. Here we defined the land-to-river and upstream-to-downstream bioinformation transportation, which is driven by watershed ecosystem processes, as the watershed biological information flow (WBIF). WBIF integrates the ecological processes of eDNA, including the origin, state, transport, and fate of eDNA (Barnes & Turner, 2016; Jo et al. 2019; Shogren et al., 2017; Cristescu and Hebert 2018; Tillotson et al. 2018). The transportation effectiveness of WBIF mainly relies on the transport capacity, degradation rate, and environmental filtration of WBIF (Barnes & Turner, 2016; Jo et al. 2017; Shogren et al., 2017; Tillotson et al. 2018). The transport capacity of WBIF mainly depends on erosion and runoff (Shogren et al., 2017; Fremier et al. 2019; Seymour, 2019); the degradation rate of WBIF mainly depends on environmental features (Barnes & Turner, 2016; Eichmiller et al. 2016; Nukazawa, et al., 2018; Shogren et al. 2018; Tillotson et al. 2018); the environmental filtration of WBIF mainly depends on the environmental change of restricting organisms; and all of these factors are related to the season and weather conditions (Nukazawa, et al., 2018). Therefore, we hypothesized that the monitoring effectiveness of riverine water eDNA would vary with the season and weather conditions. Moreover, because of species-specific eDNA degradation rates (Deiner & Altermatt, 2014), taxonomy-specific eDNA degradation rates (Barnes, et al., 2014), and form-specific eDNA degradation rates (van Bochove et al., 2020), we hypothesized that the monitoring effectiveness of riverine water eDNA would vary with taxonomic communities.

To verify the approach of using riverine water eDNA to simultaneously monitor aquatic and terrestrial biodiversity, we assessed its monitoring effectiveness, which was approximately indicated by the transportation effectiveness of land-to-river and upstream-to-downstream WBIF. In our case study that was conducted in a watershed on the Qinghai-Tibet Plateau, we estimated the monitoring effectiveness indicated by three taxonomic communities in three seasons and weather conditions. Our objectives were threefold: (1) identify the optimal holistic biodiversity monitoring season and weather conditions, (2) identify the effectiveness variation for monitoring different taxonomic communities, (3) assess the feasibility of using riverine water eDNA to simultaneously monitor aquatic and riparian biodiversity. Our work tested the approach of assessing the monitoring effectiveness of riverine water eDNA, and identified the viability of using riverine water eDNA to simultaneously monitor aquatic and terrestrial biodiversity on the Qinghai-Tibet Plateau. As the bioinformation in WBIF includes all taxonomic communities, all taxonomic communities information could be monitored using riverine water eDNA, although monitoring effectiveness variability exists among different taxonomic communities. We encourage more studies on monitoring effectiveness for each taxonomic community in other watersheds with different environmental conditions. In future ecological research, biodiversity conservation, and ecosystem management, riverine water eDNA may be a general material for routine watershed biodiversity monitoring and assessment.

## 2 Methods

### 2.1 Study Area

The Shaliu River basin (37°10′-37°52′ N, 100°17′-99°32′ E), as a sub-basin of the Qinghai Lake basin, is located 3196 m above sea level on the Qinghai-Tibet Plateau (Fig. 1). The Shaliu River is 106 km long, with a catchment area of 1320 km^2^. Grassland is the main land cover type, accounting for more than 90% of the watershed area. Less than 5% of the watershed area has been seriously changed by human activity, such as transformation into cultivated land and building land^1^. Due to its simple ecosystem assemblages and weak disturbance by human activity, the Shaliu River basin is a natural simplified model for investigating the effectiveness of monitoring aquatic and terrestrial biodiversity information using riverine water eDNA.

**Fig. 1.**
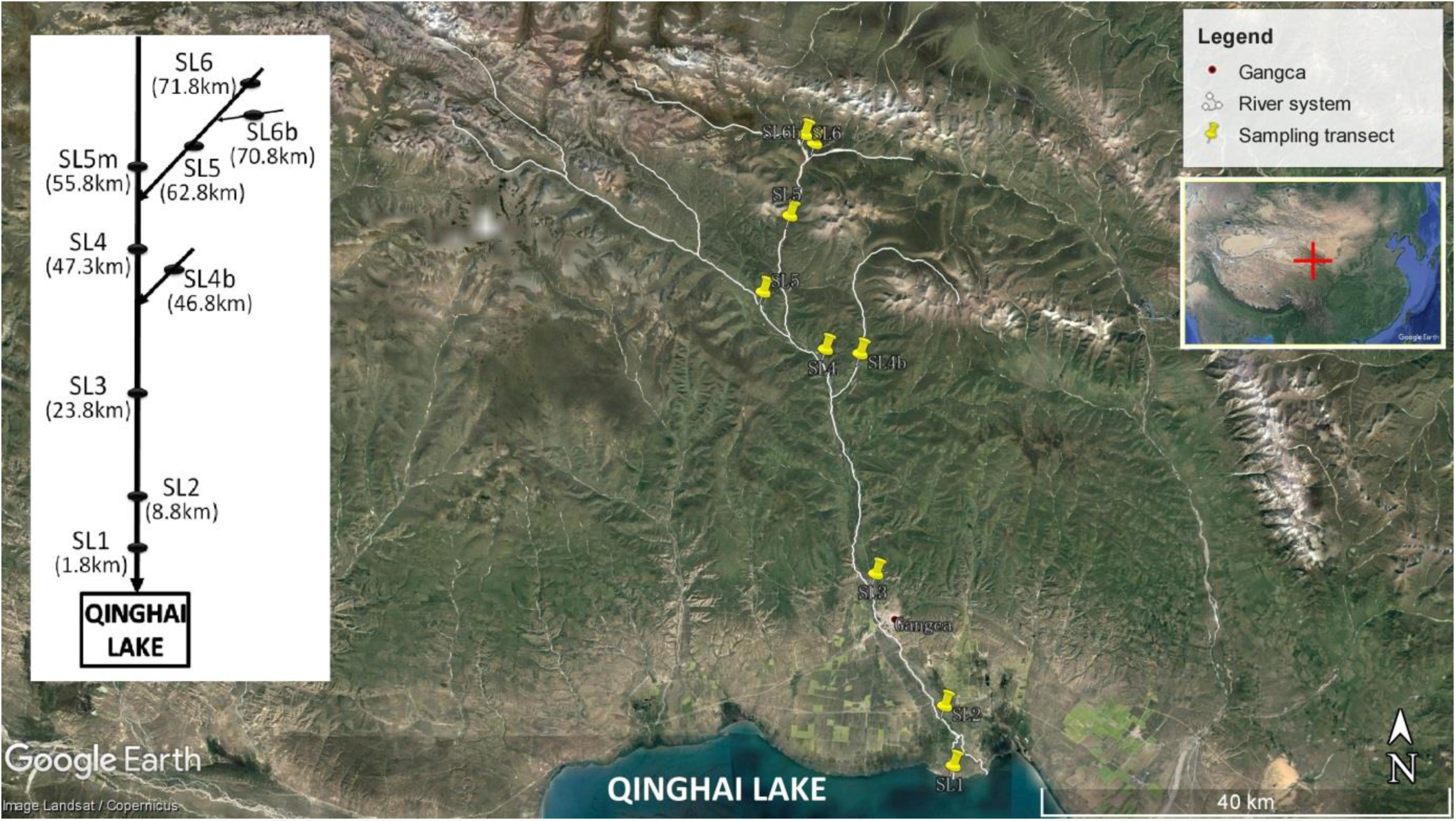
Sampling transects. SL1 denotes the first sampling transect on the Shaliu River. The distances labeled in parentheses under the tags of sampling transects denote the distances from the estuary to the sampling transects, such as SL1 (1.8 km), which means the distance from the estuary to SL1 is 1.8 km.

### 2.2 Sampling and Sequencing

To assess the monitoring effectiveness (approximately indicated by the WBIF transportation effectiveness), we compared the biodiversity information detected in adjacent upstream-to-downstream samples and that detected in adjacent riparian-to-water samples. To identify the seasonal variation of monitoring effectiveness, we sampled three times respectively in spring, summer, and autumn. To identify the taxonomic variation of monitoring effectiveness, we analyzed three taxonomic communities using the metabarcoding of the 16S rRNA, ITS, and mitochondrial CO1 genes (Collins et al., 2019; Heeger et al., 2019; Giebner et al., 2020). To obtain the eDNA of all taxonomic communities, we selected the finest filter membrane (0.2 μm) to filter riverine water samples (Eichmiller, et al., 2016; Li, et al., 2018). Because keeping the samples cool could reduce the rate of eDNA decay and is a convenient and efficient method for conserving eDNA samples (Sales, et al., 2019), we kept our eDNA samples in an ice bath or in a dry ice bath when they were transported and in an ultra-low temperature freezer when they were stored.

On April 8 and 9, June 25, and 26 and September 19 and 20, 2019, we collected eDNA samples (spring group, summer group, and autumn group), including 27 soil eDNA samples and 27 water eDNA samples, from 9 transects (including riverine sampling sites and riparian sampling sites) of the Shaliu River (Fig. 1). The weather and hydrological conditions of each group are summarized in Table 1.

**Table 1.** The weather and hydrological conditions at each sampling time.

|  | Sampling time | Air temperature | Water temperature | Discharge | Flow velocity | Weather conditions |
| --- | --- | --- | --- | --- | --- | --- |
| Spring group | 2019.4.8-9 | -6-8 °C | -0.5-0.7 °C | 1.8-3.9 m<sup>3</sup>/s | 0.63-1.04 m/s | frozen days (†)<br>frozen days (‡) |
| Summer group | 2019.6.25-26 | 7-17 °C | 4.3-16.4 °C | 29.9-45.5 m<sup>3</sup>/s | 0.84-2.03 m/s | sunny day (†)<br>light rain day (‡) |
| Autumn group | 2019.9.19-20 | 0-10 °C | 0.2-8.8 °C | 5.7-12.8 m<sup>3</sup>/s | 0.57-0.88 m/s | light rain day (†)<br>cloudy day (‡) |

(†) Collecting 4 samples (2 water samples and 2 soil samples) from transects of SL1 and SL2 (two downstream transects) along the downstream-to-upstream direction.
(‡) Collecting 14 samples (7 water samples and 7 soil samples) from transects of SL3, SL4, SL4b, SL5m, SL5, SL6 and SL6b (seven upstream transects) along the downstream-to-upstream direction.

A 5-mL soil eDNA sample (actual surface soil) was collected using a 5-mL sterilized centrifuge tube from the riparian site (5 m from the river) of each transect. A 1.5-L surface water sample (actual riverine water) was collected using a 1.5-L sterilized bottle (rinsed three times with sampling water) from the river site of each transect. Field samples were transported in an ice bath to the laboratory of the Rescue and Rehabilitation Center of Naked Carps of Qinghai Lake at 0°C. Water samples were filtered using 0.2-μm membrane filters (JinTeng, Tianjin, PRC) to obtain the eDNA sample in the laboratory (every step follows the general rule of molecular ecology experiment to control contamination and using bleach to wash the experimental apparatus); then, the filter membranes of each water eDNA sample were placed in a 50-mL sterilized centrifuge tube. The tubes with eDNA samples were frozen at −20°C, transported at −20°C (in a dry ice bath), and stored at −80°C (in an ultra-low temperature freezer) until DNA extraction. More details are provided in Table 2 and Supplementary Material 1.

**Table 2.**
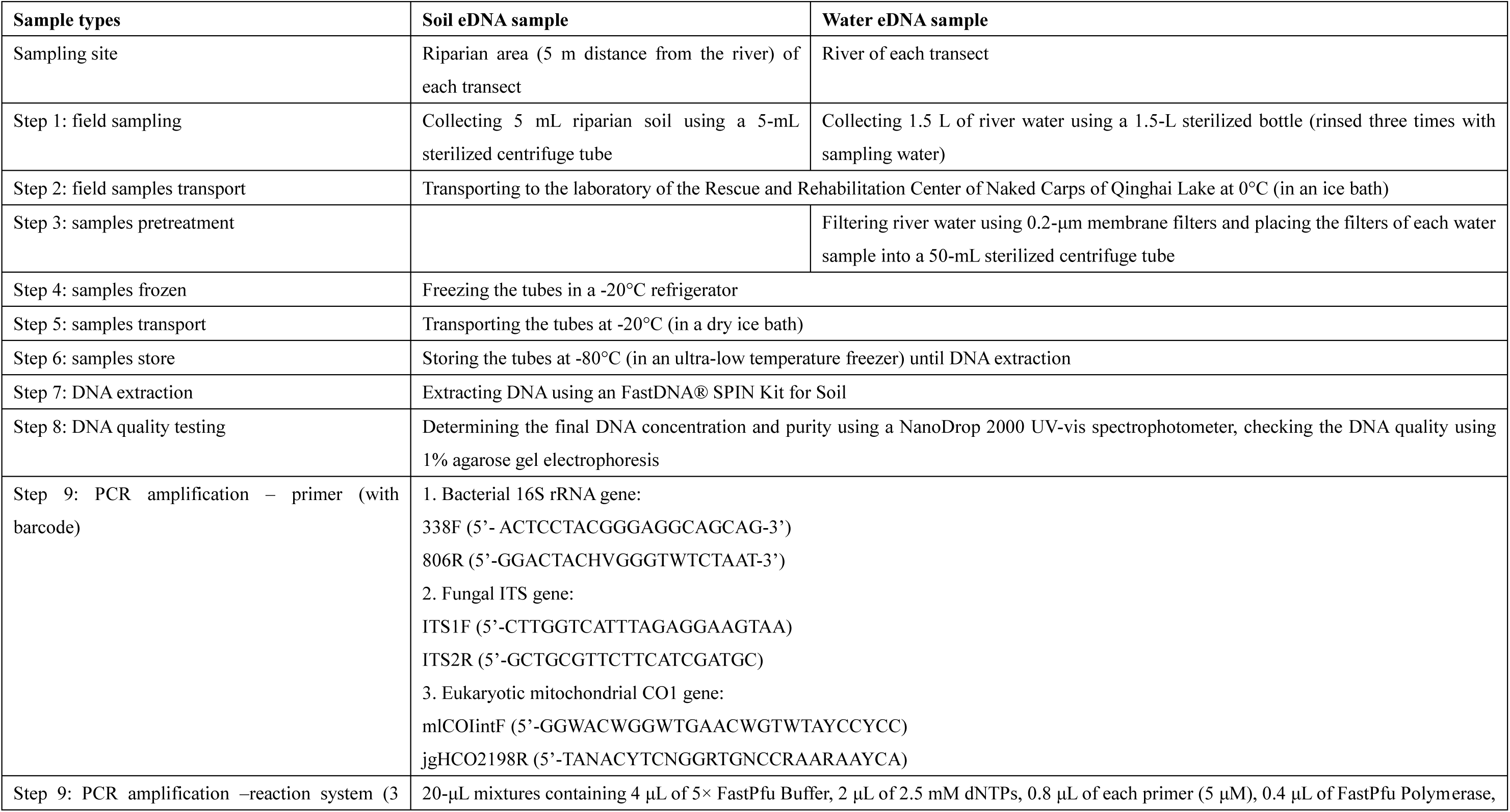
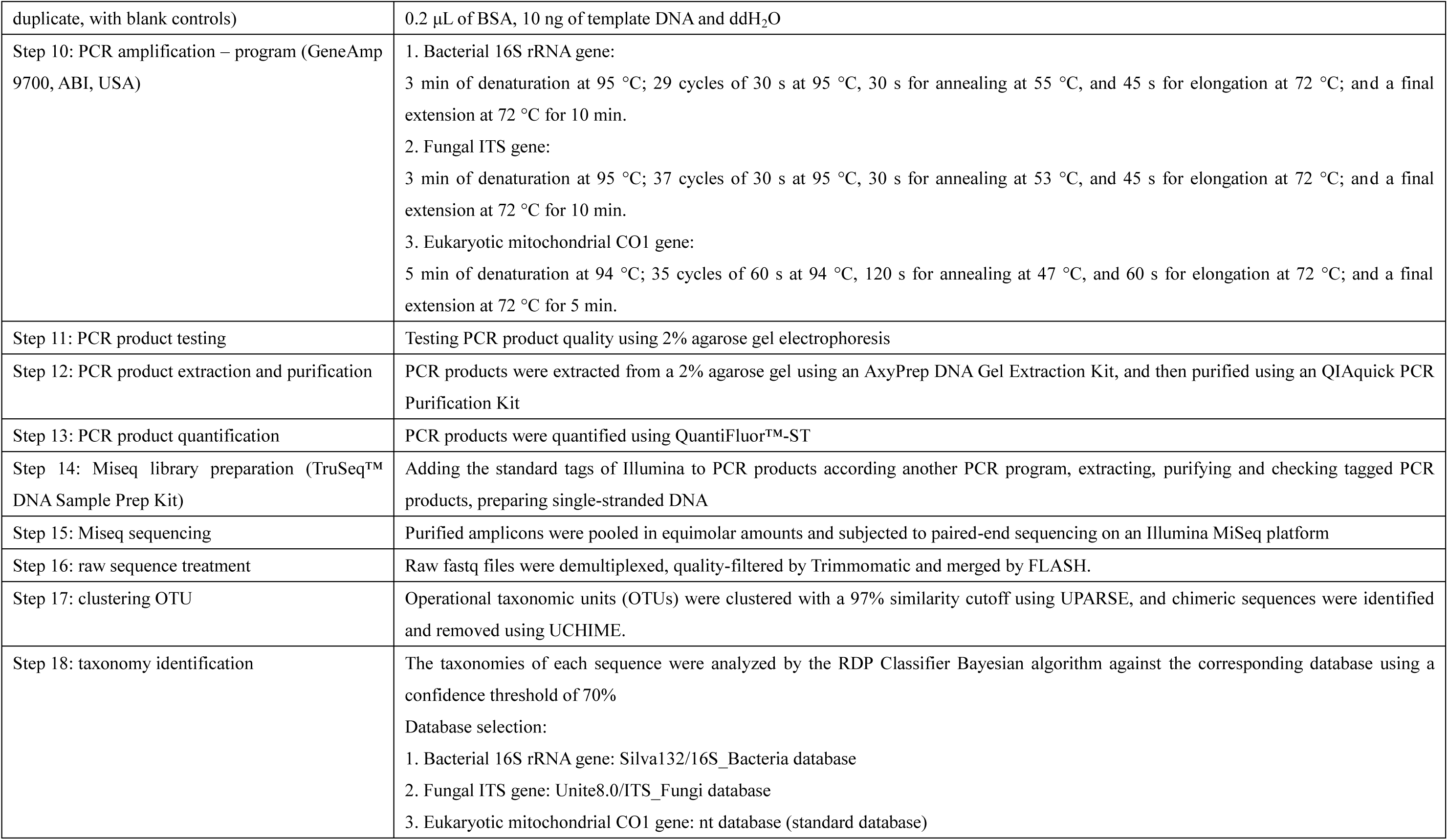
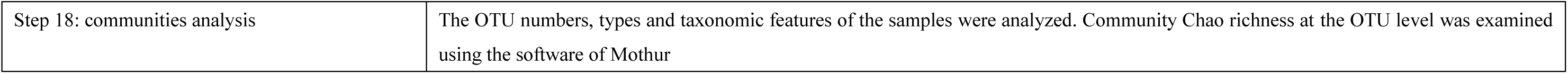
the steps of sampling and sequencing.

As the extraction of eDNA (Hermans et al., 2018; Armbrecht et al., 2020), metabarcoding selection (Collins et al., 2019; Heeger et al., 2019; Giebner et al., 2020), amplification approach, and sequencing (Nichols et al., 2018) impact the results of eDNA monitoring, a consistent DNA extraction method, primer usage, and amplification approach should be used for comparisons among samples (Dopheide et al., 2019; Giebner et al., 2020; Nicholson et al., 2020). Commercial eDNA labs can help (Ravindran, 2019), in which all approaches (including eDNA extraction, primer synthesis, amplification approach, sequencing, and contamination control, among others) could be standard. We send them the samples and select the primers, and then they give us the results — a set of sequences — and provide an interactive platform to analyze the data.

In our work, samples were processed by Shanghai Majorbio Bio-pharm Technology Co., Ltd (Shanghai, China). As long DNA fragments show a higher decay rate than short fragments (Jo, et al., 2017; Shogren, et al., 2018), short fragments better reflect community richness than long fragments (Wei, et al., 2018; Jo, et al., 2019). Thus, we restricted the amplified fragment length to 300-500 bp. To identify the taxonomic communities, we selected the primers 338F/806R, ITS1F/ ITS2R, and mlCOIintF/ jgHCO2198R to indicate bacteria, fungi, and metazoan, respectively (Collins et al., 2019; Heeger et al., 2019; Giebner et al., 2020). The details on regarding the DNA extraction, PCR amplification, fluorescence quantitation, library preparation, and Illumina Miseq sequencing are provided in Table 2 and Supplementary Material 1.

We analyzed the types of operational taxonomic unit (OTU), the sequence number of each OTU, and the taxonomic features of each sample and examined the community richness (Chao richness index at the OTU level) to reveal the variation among the three groups using Mothur software. The raw data have been deposited in the China National GeneBank Sequence Archive (CNSA, https://db.cngb.org/cnsa/) of the China National GeneBank database (CNGBdb) under accession number CNP0001046. The data were analyzed on the free online Majorbio Cloud Platform (www.majorbio.com).

### 2.3 WBIF Analysis

The WBIF (including land-to-river and upstream-to-downstream WBIF) of each group was assessed to reveal the effectiveness of monitoring the upstream and riparian biodiversity using riverine water eDNA. In the current WBIF analysis, all statistics used the types of OTUs and species in each sample. The processing approach was simply described as follows (indicated by the OTU type).

The transportation effectiveness of land-to-river and upstream-to-downstream WBIF could be estimated by comparing the OTU assemblages between the adjacent soil eDNA sample and water eDNA sample and by comparing the OTU assemblages between adjacent (upstream-downstream) water eDNA samples. The transportation effectiveness of WBIF was indicated by the proportion of input OTU types (i.e., the common types between the source site sample and the pool site sample) to output OTU types (the total types of source site sample) (Eq. 1).

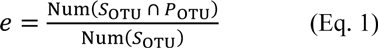

where *e* denotes the transportation effectiveness of WBIF; *S_OTU_* denotes the OTU assemblage of the source site sample (i.e., the adjacent soil eDNA sample in the land-river WBIF or the adjacent upstream water eDNA sample in the upstream-downstream WBIF); and *P_OTU_* denotes the OTU assemblage of the pool site sample (i.e., the adjacent water eDNA sample in the land-river WBIF or the adjacent downstream water eDNA sample in the upstream-downstream WBIF).

As the transportation effectiveness of WBIF relied on the transport capacity, degradation rate, and environmental filtration and the distance of the land-to-river WBIF was less than 5 m in this case study, the transportation effectiveness of the land-to-river WBIF was assumed to be constructed by transport capacity and environmental filtration (no degradation rate). The transportation effectiveness of the land-to-river WBIF could be indicated by the proportion of the common types shared between adjacent soil eDNA samples and water eDNA samples to the total types of soil eDNA samples (Eq. 1). The transport capacity of the land-to-river WBIF could be indicated by the proportion of the common types shared between adjacent soil eDNA samples and water eDNA samples to the common types shared between the soil eDNA sample and all water eDNA samples in the corresponding group (Eq. 2). The environmental filtration of the land-to-river WBIF could be indicated by the proportion of the types included in the soil eDNA sample, but not in any water eDNA sample to the total types in the soil eDNA sample (Eq. 3).

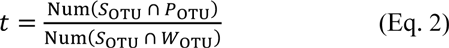

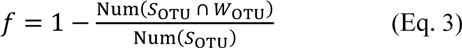

where *t* denotes the transport capacity; *f* denotes the environmental filtration; *S_OTU_* denotes the OTU assemblage of the source site sample (i.e., the soil eDNA sample); and *W_OTU_* denotes the OTU assemblage of all water eDNA samples.

WBIF included the effective WBIF (i.e., the flow or migration of living organisms) and noneffective WBIF (i.e., the flow of the bioinformation labeling the biological material without life activity and fertility (dead bioinformation)). The transportation effectiveness of upstream-to-downstream WBIF was determined by the different features of effective WBIF and noneffective WBIF. The effective WBIF was impacted by transport capacity and environmental filtration. The noneffective WBIF was impacted by transport capacity and degradation rate. We established the following presuppositions: (1) the transport capacity was consistent in a defined runoff condition of a definite season and weather condition; (2) the proportion of noneffective WBIF at each site was consistent; (3) the noneffective WBIF degraded over time (i.e., distance) in a logistic manner; and (4) the environmental filtration was consistent in a definite environmental change. These four presuppositions did not exactly describe the complex facts, but provided a possibility of constructing a model to approximately deal with the complex facts. The transportation effectiveness of the upstream-to-downstream WBIF could be constructed by the transport capacity of WBIF, the environmental filtration of the effective WBIF and the degradation rate of the noneffective WBIF. It could be described by an equation (Eq. 4), in which the transportation effectiveness was the function of runoff distance, and then the transport capacity, environmental filtration and degradation rate were parameters that could be estimated according to the sets of transportation effectiveness and runoff distance. In practice, as WBIF are impacted by varied influencing factors at any site and time, the analytical solution of parameters in Eq. 4 is impossible. Therefore, we suggested that Eq. 4 could be programming-solved according to the evolutionary algorithm in Microsoft Excel. As there were only approximate solutions of parameters in Eq. 4, we suggested getting several sets (such as 30 sets) of approximate solutions and then taking a statistics for each parameter.

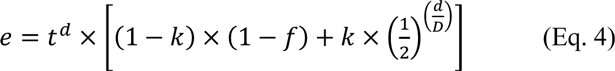

where *e* denotes the transportation effectiveness of WBIF; *t* denotes the transport capacity; *d* denotes the distance of WBIF; *k* denotes the proportion of the noneffective WBIF; *f* denotes the environmental filtration; and *D* denotes the half-life distance.

In this case study, we first sequenced the bacterial 16S rRNA gene of the eDNA samples from three seasonal groups and assessed the transportation effectiveness of WBIF (indicated by microbial OTUs) at three seasons. Second, we selected the eDNA samples of the seasonal group that showed the highest WBIF transportation effectiveness, sequenced their fungal ITS gene and eukaryotic mitochondrial CO1 gene and assessed the WBIF transportation effectiveness indicated by three taxonomic communities (bacteria, fungi and metazoan) at both the OTU level and species level.

## 3 Results

### 3.1 WBIF Features of the Three Seasonal Groups

A total of 10,602, 13,766, and 16,500 bacterial OTU types were detected from the samples (including 9 water samples and 9 soil samples) of the spring, summer, and autumn group, respectively (Fig. 2 and Table S1). The total types of OTUs that were detected from the soil eDNA samples were similar among the seasons (Figs. 2,3). The total types of OTUs that were detected from the water eDNA samples were richest in autumn (Figs 2,3). The common types of OTUs that were shared between the soil eDNA and water eDNA samples accounted for 36.30%, 71.98%, and 67.58% of the total OTU types that were detected in the soil eDNA samples in the spring, summer, and autumn group, respectively (Fig. 3).

**Fig. 2.**
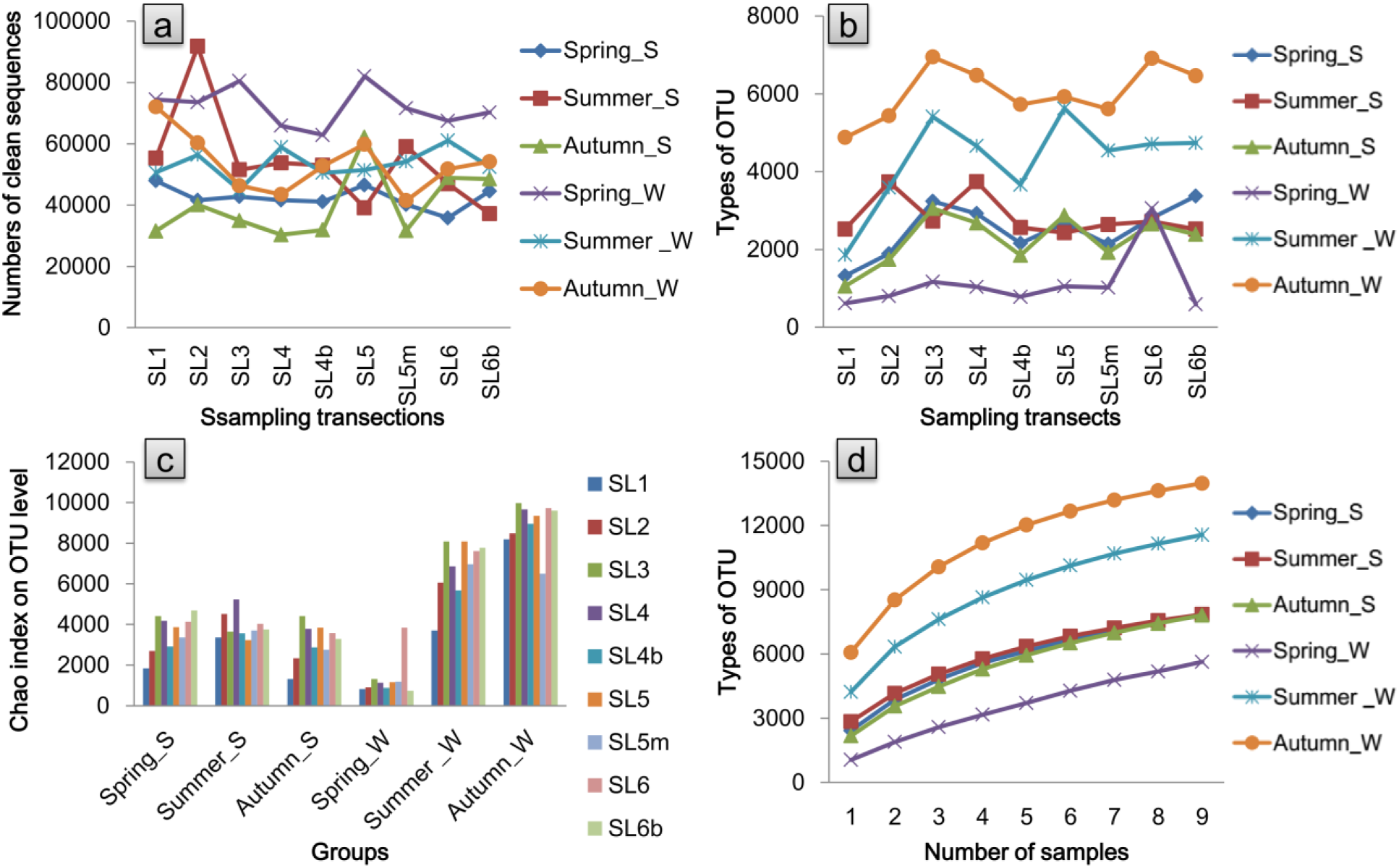
Biological information features of the samples: numbers of clean sequences in each sample (a), types of OTUs in each sample (b), community richness of each sample at the OTU level (c) and species accumulation curves at the OTU level (d) Spring_S denotes the soil eDNA samples that were sampled during April 2019; Spring_W denotes the water eDNA samples that were sampled during April 2019; Summer_S denotes the soil eDNA samples that were sampled during June 2019; Summer _W denotes the water eDNA samples that were sampled during June 2019; Autumn_S denotes the soil eDNA samples that were sampled during September 2019; Autumn_W denotes the water eDNA samples that were sampled during September 2019.

**Fig. 3.**
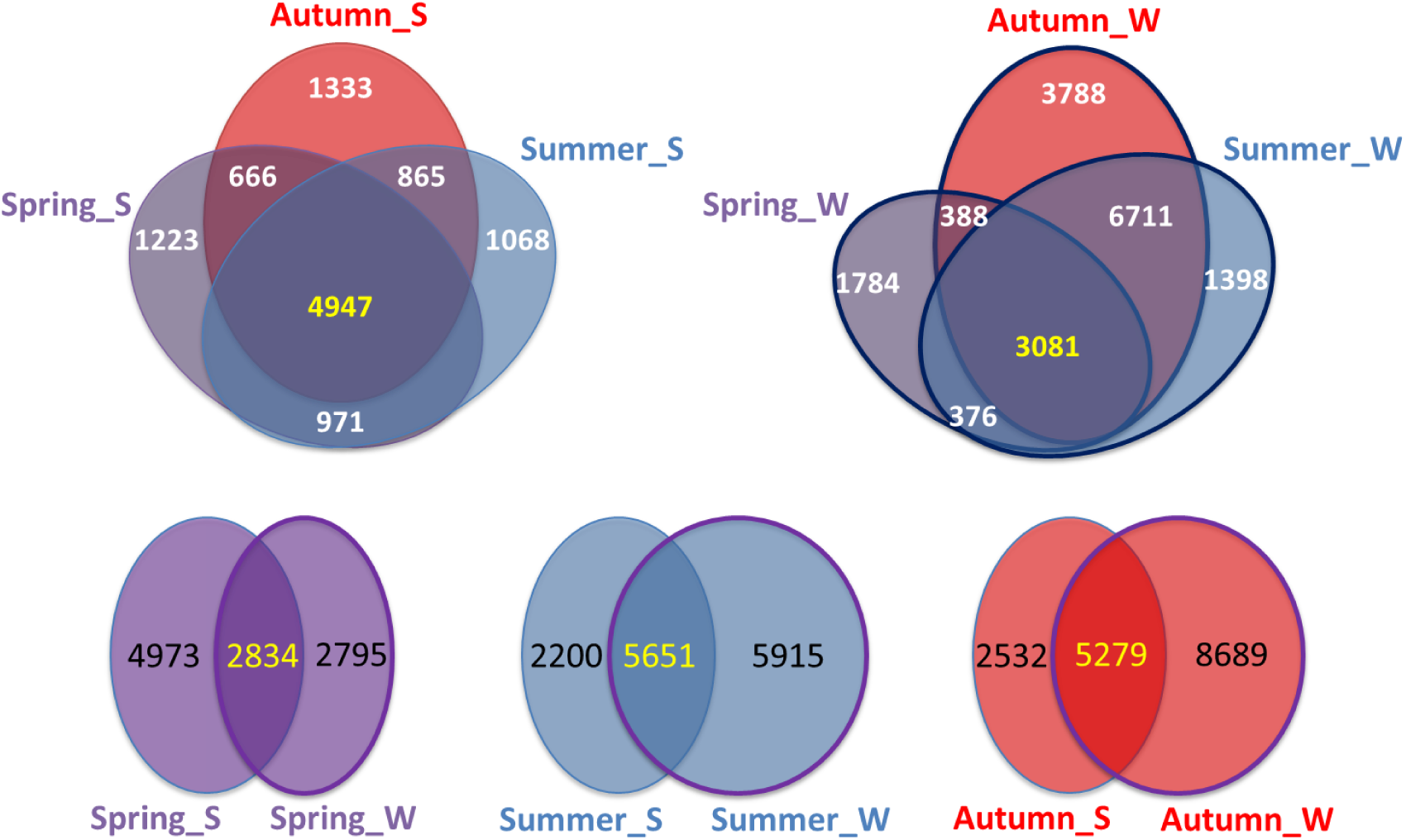
OTU types in riparian soil samples and riverine water samples shared by the three groups. Spring_S denotes the soil eDNA samples that were sampled during April 2019; Spring_W denotes the water eDNA samples that were sampled during April 2019; Summer_S denotes the soil eDNA samples that were sampled during June 2019; Summer _W denotes the water eDNA samples that were sampled during June 2019; Autumn_S denotes the soil eDNA samples that were sampled during September 2019; Autumn_W denotes the water eDNA samples that were sampled during September 2019.

The transportation effectiveness of WBIF indicated by bacterial OTUs from the riparian sampling site to adjacent riverine sampling site was 16.62%, 62.76%, and 48.09% on spring frozen, summer rainy, and autumn cloudy days, respectively, among which there was the highest transport capacity and the lowest environmental filtration on the summer rainy day (Tables 3,S2). The transportation effectiveness of WBIF indicated by bacterial OTUs from upstream to downstream was 75.86%, 97.41%, and 96.07% per km on spring frozen, summer rainy, and autumn cloudy days, respectively (Tables 4,S3), among which the transport capacity was more than 99% in all three seasons and the least noneffective WBIF (dead bioinformation) occurred; the longest half-life distance of the noneffective WBIF occurred on the summer rainy day (Table 4).

**Table 3.** Seasonal variation of transport capacity, environmental filtration, and transportation effectiveness of watershed biological information flow (WBIF) from the riparian sampling site to adjacent riverine water sampling site in three seasons indicated by bacterial OTUs.

| Seasonal group | Weather condition | Transport capacity | Environmental filtration | Transportation effectiveness |
| --- | --- | --- | --- | --- |
| Spring group | Frozen days | 0.268791 ±0.202388 | 0.385443 ±0.029320 | 0.166152 ±0.125394 |
| Summer group | Rainy days | 0.684876 ±0.091302 | 0.083816 ±0.020574 | 0.627643 ±0.087327 |
| Autumn group | Cloudy days | 0.573579 ±0.052897 | 0.161800 ±0.045075 | 0.480933 ±0.052179 |

The spring group was sampled during April 2019; the summer group was sampled during June 2019; the autumn group was sampled during September 2019. Statistics for the spring group are based on 8 sampling transects except estuary (SL1); statistics for the summer and autumn groups are based on 7 sampling transects except two downstream transects (SL1 and SL2).

**Table 4.** Seasonal variation of transport capacity, proportion of noneffective WBIF, half-life distance of the noneffective WBIF, and transportation effectiveness of watershed biological information flow (WBIF) from the upstream to downstream regions indicated by bacterial OTUs.

| Seasonal group | Weather condition | Transport capacity (per km) | Proportion of noneffective WBIF | Half-life distance of the noneffective WBIF | Transportation effectiveness (per km) | Environmental filtration from rain point to sunny point | Environmental filtration from freshwater to saline-water |
| --- | --- | --- | --- | --- | --- | --- | --- |
| **Spring group** | Frozen days | 0.999706±0.000305 | 0.668465±0.003435 | 1.548987±0.126870 | 0.758618±0.000304 | / | 0.160427±0.008244 |
| **Summer group** | Rainy days | 0.994245±0.000941 | 0.434635±0.041681 | 14.52338±1.440539 | 0.974105±0.000926 | 0.005687±0.005450 | 0.544164±0.010042 |
| **Autumn group** | Cloudy days | 0.992250±0.001452 | 0.493504±0.041043 | 10.398112±0.711122 | 0.960671±0.001415 | / | 0.128718±0.017062 |

The spring group was sampled during April 2019; the summer group was sampled during June 2019; the autumn group was sampled during September 2019.

### 3.2 WBIF Features of the Three Taxonomic Groups

A total of 13,766, 7098, and 17,316 kinds of OTUs and 3532, 1032, and 6836 kinds of species were detected among the 18 summer samples, as indicated by the 16S rRNA gene, ITS gene, and CO1 gene, respectively (Fig. 4 and Table S4). The types of OTUs and species detected in the water eDNA samples were generally higher than in the soil eDNA samples for all three taxonomic communities (Fig. 4). The common OTUs and species shared between the soil and water eDNA samples accounted for 71.98% and 87.95%, 60.40% and 76.18%, and 37.93% and 53.52% of the total types of OTUs and species in the bacterial, fungal and eukaryotic group, respectively.

**Fig. 4.**
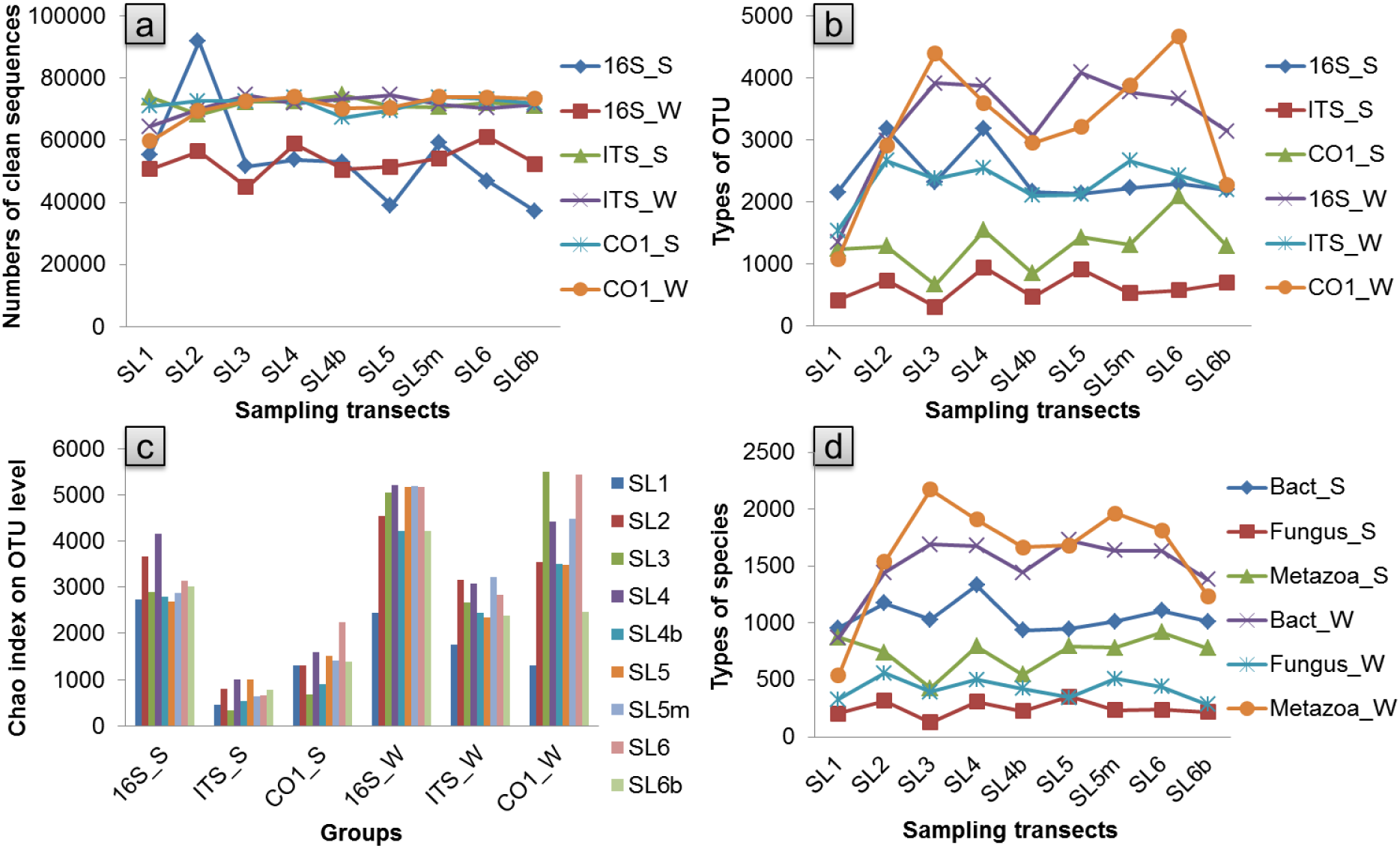
Biological information features of the samples: numbers of clean sequences in each sample (a), types of OTUs in each sample (b), community richness of each sample at the OTU level (c), and types of species in each sample (d) 16S_S denotes the soil eDNA samples that were sequenced using the bacterial 16S rRNA gene; ITS_S denotes the soil eDNA samples that were sequenced using the fungal ITS gene; CO1_W denotes the water eDNA samples that were sequenced using the eukaryotic mitochondrial CO1 gene. Bac_S denotes the bacterial group detected in the soil eDNA samples; Fungus_S denotes the fungal group detected in the soil eDNA samples; and Metazoa_W denotes the metazoan group detected in the water eDNA samples.

The transportation effectiveness of the bacterial, fungal, and eukaryotic WBIF from the riparian sampling site to the adjacent riverine sampling site was 62.76%, 44.79%, and 22.64% at the OTU level, respectively, and 80.75%, 65.62%, and 43.38% at the species level, respectively, among which both the transport capacity and environmental filtration significantly declined with the bacterial, fungal, and eukaryotic communities (Tables 5,S5,S6). The transportation effectiveness of bacterial, fungal and eukaryotic WBIF from upstream to downstream was 97.41%, 92.64% and 89.83% per km at the OTU level, and 98.69%, 95.71% and 92.41% per km at the species level, respectively, among which the noneffective WBIF decreased with the bacterial, fungal, and eukaryotic communities (Tables 6,S7,S8), and the half-life distance of the noneffective WBIF was 14.52, 4.93, and 4.07 km at the OTU level and 17.82, 5.96, and 5.02 km at the species level for the bacterial, fungal, and eukaryotic groups, respectively (Table 6).

**Table 5.** Transport capacity, environmental filtration, and transportation effectiveness of watershed biological information flow (WBIF) from the riparian sampling site to the adjacent riverine water sampling site on summer rainy days, indicated by three taxonomic groups.

| **Taxonomic group** | **Taxonomic level** | **Transport capacity** | **Environmental filtration** | **Transportation effectiveness** |
| --- | --- | --- | --- | --- |
| **Bacteria (detected by the 16S rRNA gene)** | OTU level | 0.684876±0.091302 | 0.083816±0.020574 | 0.627643±0.087327 |
|  | species level | 0.829912±0.066079 | 0.027020±0.007048 | 0.807461±0.064521 |
| **Fungi (detected by the ITS gene)** | OTU level | 0.600756±0.102865 | 0.258922±0.054794 | 0.447896±0.095670 |
|  | species level | 0.738975±0.100006 | 0.113469±0.016910 | 0.656191±0.097099 |
| **Metazoan (detected by the CO1 gene)** | OTU level | 0.440871±0.124206 | 0.485954±0.061102 | 0.226403±0.071669 |
|  | species level | 0.604263±0.092950 | 0.281177±0.028991 | 0.433842±0.066684 |

Bacteria (detected by the 16S rRNA gene), fungi (detected by the ITS gene), and metazoan (detected by the CO1 gene) indicate the groups of bacteria (detected by the 16S rRNA gene), fungi (detected by the ITS gene), and metazoan (detected by the CO1 gene), respectively. Statistics in all groups are based on 7 sampling transects excepted two downstream transects (SL1 and SL2).

**Table 6.** Transport capacity, proportion of noneffective WBIF, half-life distance of the noneffective WBIF, and transportation effectiveness of watershed biological information flow (WBIF) from the upstream to downstream regions on summer rainy days, indicated by three taxonomic groups at the OTU and species levels estimated by programming-solved according to the evolutionary algorithm.

| Taxonomic group | Taxonomic level | Transport capacity (per km) | Proportion of noneffective WBIF | Half-life distance of the noneffective WBIF | Transportation effectiveness (per km) | Environmental filtration from rain point to sunny point | Environmental filtration from freshwater to saline-water |
| --- | --- | --- | --- | --- | --- | --- | --- |
| **Bacteria (detected by the 16S rRNA gene)** | **OTU level** | 0.994245±0.000941 | 0.434635±0.041681 | 14.52338±1.440539 | 0.974105±0.000926 | 0.005687±0.005450 | 0.544164±0.010042 |
|  | **species level** | 0.998188±0.000121 | 0.296484±0.010590 | 17.82057±1.215028 | 0.986898±0.000121 | 0.051209±0.005337 | 0.460245±0.001469 |
| **Fungi (detected by the ITS gene)** | **OTU level** | 0.995550±0.000680 | 0.529290±0.016749 | 4.925445±0.353730 | 0.926377±0.000670 | 0.003482±0.002886 | 0.338354±0.003866 |
|  | **species level** | 0.999484±0.000244 | 0.386710±0.008333 | 5.961259±0.264864 | 0.957057±0.000242 | 0.000541±0.000258 | 0.224685±0.001239 |
| **Metazoan (detected by the CO1 gene)** | **OTU level** | 0.989275±0.000923 | 0.587740±0.019079 | 4.073058±0.362046 | 0.898288±0.000908 | 0.007897±0.006958 | 0.716408±0.003182 |
|  | **species level** | 0.992862±0.000724 | 0.537202±0.016816 | 5.018684±0.317762 | 0.924058±0.000713 | 0.005337±0.002702 | 0.607287±0.002642 |

Bacteria (detected by the 16S rRNA gene), fungi (detected by the ITS gene), and metazoan (detected by the CO1 gene) indicate the groups of bacteria (detected by the 16S rRNA gene), fungi (detected by the ITS gene), and metazoan (detected by the CO1 gene), respectively.

## 4 Discussion

### 4.1 The optimal holistic biodiversity monitoring season and weather conditions

This study showed that summer and autumn were the optimal seasons and rainy days were the optimal weather condition for using riverine water eDNA to simultaneously monitor aquatic and terrestrial holistic biodiversity. Our results indicated that the season did not impact the soil bacterial community richness, but it did impact the water bacterial community richness (Fig. 2). The water bacterial community composition was richest in autumn followed by summer (Fig. 2,3). Thus, the richest biodiversity was found in autumn followed by summer. Considering the biodiversity in soil and water, autumn and summer were the optimal holistic biodiversity monitoring seasons.

Our results indicated that the transportation effectiveness of riparian-to-water WBIF in spring was significantly lower than in summer and autumn (Tables 3,S2). The transportation effectiveness of riparian-to-water WBIF in the summer group on a sunny day (transect SL2) was significantly lower than that on a rainy day (Tables 3,S2). The transportation effectiveness of riparian-to-water WBIF in the autumn group on a cloudy day was also significantly lower than on a rainy day (transect SL2) (Tables 3,S2). There were no significant differences between the transportation effectiveness of riparian-to-water WBIF in the summer group on a sunny day (transect SL2) and in the autumn group on a cloudy day (Tables 3,S2).

There were no significant differences between the transportation effectiveness of riparian-to-water WBIF in the summer group on a rainy day and in the autumn group on a rainy day (transect SL2) (Tables 3,S2). Thus, there was a similar transportation effectiveness of riparian-to-water WBIF in summer and autumn, and both were higher than in spring. The rain (although light rain, less than 10 mm) increased the transportation effectiveness of riparian-to-water WBIF in both summer and autumn. Considering the monitoring effectiveness of adjacent water eDNA to riparian bacterial communities (indicated by the transportation effectiveness of riparian-to-water WBIF), summer and autumn were the optimal holistic biodiversity monitoring seasons, and rainy days represented the optimal holistic biodiversity monitoring weather condition.

Our results indicated that the transportation effectiveness of upstream-to-downstream WBIF in spring was significantly lower than in summer and autumn (Tables 4,S3). Although the transportation effectiveness of upstream-to-downstream WBIF on summer rainy days was significantly higher than on autumn cloudy days, considering the impact of rain (environmental filtration from rain point to a sunny point), there were no significant differences between the transportation effectiveness of upstream-to-downstream WBIF in summer and autumn (Table 4). Thus, there was a similar transportation effectiveness of upstream-to-downstream WBIF in summer and autumn, and both were higher than in spring. The rain (leading to turbid river water) increased the transportation effectiveness of upstream-to-downstream WBIF, although it was slight. Considering the monitoring effectiveness of adjacent water eDNA to upstream bacterial communities (indicated by the transportation effectiveness of upstream-to-downstream WBIF), summer and autumn were the optimal holistic biodiversity monitoring seasons, and rainy days represented the optimal holistic biodiversity monitoring weather condition.

Previous studies have pointed out that, driven by resource availability and temperature, there are significant seasonal shifts in the structures and functions of soil microbial communities (Koranda et al., 2013; Xu et al., 2018). Similarly, in our work, the soil bacterial community composition also shifted in three seasons (Fig. 3). However, there were no significant differences in the soil bacterial community richness among the three seasons (Fig. 2). In summer and autumn, along with the high microbial activity impacted by temperature (Lin et al., 2016), the airborne microbiota, animal symbiotic microorganisms, and terrestrial soil microbes with input into the river system enriched the water microbial communities (Deiner et al., 2016; Gusareva et al. 2019; Matsuoka et al., 2019). Especially, according to the surface runoff caused by rain, more terrestrial soil microbes were input into the river system and more microbes were preserved in soil aggregates (Wilpiszeski et al. 2019). In our work, the richest water bacterial communities were found in autumn and summer (Fig. 2,3). The highest proportion of riparian soil bacteria was input into the river system and preserved in water in summer and autumn, and the rain promoted this phenomenon (Fig. 3 and Tables 3,S2).

The monitoring of upstream biodiversity using downstream water eDNA depends on the biodiversity information transport from upstream to downstream (Deiner & Altermatt,2014; Sansom & Sassoubre,2017; Pont, et al., 2018; Seymour,2019). The biodiversity information detected by water eDNA could originate from living and dead organisms (Stoeckle et al., 2016; Nukazawa et al., 2018; Tillotson et al., 2018). The detection of biodiversity information that originates from a living organism depends on the dispersal of this living organism (Pont, et al., 2018; Ravindran,2019). The detection of biodiversity information that originates from a dead organism depends on its degradation rate (Seymour et al., 2018; Jo et al., 2017; Seymour,2019). eDNA (biodiversity information originates from dead organism) degradation is impacted by many environmental conditions, such as the biochemical oxygen demand, temperature, pH, organic matter, and substrate (Barnes, et al., 2014; Eichmiller, et al., 2016; Shogren, et al., 2017; Nukazawa, et al., 2018; Seymour, et al., 2018; van Bochove, et al., 2020). eDNA degrades over time in a logistic manner (half-life time) (Barnes, et al., 2014; Jo et al., 2017; Shogren, et al., 2018; Wei et al., 2018; Seymour,2019). In other words, eDNA degrades over distance in a logistic manner (half-life distance) in lotic system. In our work, impacted by temperature (Table 1) (Lin et al., 2016), the biodiversity information originating from living organism (proportion of effective WBIF) in summer and autumn was significantly higher than in spring (Table 4). Driven by soil aggregates (Wilpiszeski et al. 2019), the proportion of effective WBIF was significantly higher on rainy than on cloudy days (Table 4). Driven by substrate absorption that was influenced by discharge (Table 1) (Shogren, et al., 2017), the half-life distance of noneffective WBIF in spring was far less than in summer and autumn (Table 4). Driven by flow velocity (Table 1), the half-life distance of noneffective WBIF was significantly farther in summer than in autumn (Table 4). Thus, using riverine water eDNA to simultaneously monitor aquatic and riparian biodiversity, summer and autumn were the optimal holistic biodiversity monitoring seasons, and rainy days were the optimal holistic biodiversity monitoring weather condition.

### 4.2 The monitoring effectiveness of each taxonomic community

Our work showed that the effectiveness of monitoring eukaryotic communities was significantly lower than monitoring bacterial communities, and the monitoring effectiveness at the OTU level was significantly lower than at the species level. Our results revealed the detection of a significantly higher proportion of bacterial communities than eukaryotic communities from riparian soil using riverine water eDNA (Table 5). A significantly higher proportion of micro eukaryotic communities (fungi) than overall eukaryotic communities (including micro- and macro-organisms) from riparian soil was detected using riverine water eDNA (Table 5). The monitoring effectiveness of using riverine water eDNA to monitor adjacent riparian site biodiversity (transportation effectiveness of riparian-to-water WBIF) at the OTU level (62.76%, 44.79%, and 22.64%, respectively, for bacteria, fungi, and metazoan on summer rainy days) was generally lower than at the species level (80.75%, 65.62%, and 43.38%, respectively, for bacteria, fungi, and metazoan on summer rainy days) (Table 5). The overall monitoring effectiveness of riverine water eDNA to riparian biodiversity on summer rainy days was 71.98%, 60.40%, and 37.93% for bacteria, fungi, and metazoan at the OTU level, and 87.95%, 76.18%, and 53.52% for bacteria, fungi, and metazoan at the species level, respectively.

Our results revealed a significantly higher downstream-to-upstream monitoring effectiveness of water eDNA (and proportion of effective WBIF) for bacterial communities than for eukaryotic communities (Table 6). A significantly higher downstream-to-upstream monitoring effectiveness of water eDNA (and proportion of effective WBIF) was found for micro eukaryotic communities (fungi) than for overall eukaryotic communities (including micro- and macro-organism) (Table 6). The downstream-to-upstream monitoring effectiveness of water eDNA at the OTU level (97.41%, 92.64%, and 89.83% per km, respectively, for bacteria, fungi, and metazoan on summer rainy days) was generally lower than at the species level (98.69%, 95.71%, and 92.41% per km, respectively, for bacteria, fungi, and metazoan on summer rainy days) (Table 6). The proportion of effective upstream-to-downstream WBIF at the OTU level (58.54%, 47.07%, and 41.23%, respectively, for bacteria, fungi, and metazoan on summer rainy days) was generally lower than at the species level (70.35%, 61.33%, and 46.28%, respectively, for bacteria, fungi, and metazoan on summer rainy days) (Table 6). The half-life distance of noneffective WBIF for bacteria was much farther than that for metazoan (Table 6). The half-life distance of noneffective WBIF on summer rainy days was respectively 14.52, 4.93, and 4.07 km for bacteria, fungi, and metazoan at the OTU level, and 17.82, 5.96, and 5.02 km for bacteria, fungi, and metazoan at the species level (Table 6).

As researchers can actually acquire information from across the taxonomic tree of life, eDNA surveys, based on metabarcoding, are becoming an increasingly powerful field-biology technique (Deiner et al., 2016; Stat et al., 2017; Wilcox, et al., 2018; Ravindran,2019; Djurhuus, et al., 2020; Pawlowski et al., 2020). However, eDNA that originates from different taxonomic groups has a different probability of being left in the environment and input into water (Deiner, et al., 2016; Harper, et al., 2019; Seeber, et al., 2019; Sales, et al., 2020). Our work indicated that more bacteria than eukaryotes and more microorganisms than macroorganisms (OUT and species types) in riparian soil left traces and were input into water (Table 5).

Moreover, previous studies have indicated that there are species-specific eDNA degradation rates (Deiner & Altermatt, 2014), taxonomy-specific eDNA degradation rates (Barnes et al., 2014), and form-specific eDNA degradation rates (van Bochove et al., 2020). Our results showed that the half-life distance of noneffective WBIF for bacteria (detected by 16s RNA gene, about 13-19 km) was much farther than that for unicellular eukaryotes (detected by ITS gene, mainly unicellular, about 5-6 km), than that for multicellular eukaryotes (detected by CO1 gene, mainly multicellular, about 4-5 km) (Table 6), which indicated that the difference between prokaryotes and eukaryotes is the most important differentiator for taxonomy-specific eDNA degradation rates. van Bochove et al. (2020) inferred that the eDNA contained inside cells is extra resilient against degradation (i.e., intracellular vs. extracellular) (van Bochove, et al., 2020). Based on our results, we inferred that eDNA contained inside bacterial cells was more resilient against degradation than that contained inside unicellular eukaryotic cells (i.e., prokaryotic cells vs. eukaryotic cells), than that contained multicellular eukaryotic cells or extracellular mitochondria (i.e., unicellular eukaryotic cells vs. multicellular eukaryotic cells or extracellular).

Mainly, a eDNA metabarcoding survey is used to detect the presence of species (Civade, et al., 2016; Wilcox, et al., 2018; Shogren, et al., 2019; Wacker, et al., 2019; Beng and Corlett 2020). Of course, this detection could be used at the OTU level (the level under species) (Giebner et al. 2020). Because of the data structure in which if one OTU type of a species is present then the species is delineated present despite the absence of several OTU type, the transportation effectiveness of WBIF at the OTU level is generally lower than at the species level (Table 5,6).

### 4.3 The viability of using riverine water eDNA to simultaneously monitoring aquatic and riparian biodiversity

Our work showed that in the optimal monitoring season and weather conditions (a summer rainy day) in a watershed on the Qinghai-Tibet Plateau (the Shaliu river basin), using riverine water eDNA, we can monitor as much as 87.95% bacterial species, 76.18% fungal species, and 53.52% eukaryotic species from riparian soil (Table 5), and as much as 98.69% bacterial species, 95.71% fungal species, and 92.41% eukaryotic species from 1 km upstream (Table 6). The half-life distance of noneffective WBIF was respectively 17.82, 5.96, and 5.02 km for bacteria, fungi, and metazoan at the species level (Table 6).

Previous studies have indicated that terrestrial organisms could be detected by using water eDNA (Rodgers & Mock,2015; Ushio, et al., 2017; Harper, et al., 2019; Seeber, et al., 2019; Sales, et al., 2020), upstream organisms could be detected by downstream water eDNA (Deiner & Altermatt,2014; Jerde, et al., 2016; Stoeckle, et al., 2016; Sansom & Sassoubre,2017; Carraro, et al., 2018; Pont, et al., 2018; Seymour,2019; Carraro, et al., 2020), and riverine water eDNA incorporates biodiversity information across terrestrial and aquatic biomes (Deiner et al., 2016; Matsuoka et al., 2019). Currently, we initially systematically quantify the capacity of monitoring riparian and aquatic biodiversity using riverine water eDNA.

Sales et al. (2020) indicated that although the detection probability of riverine water eDNA was 57% (43%-71%), 40% (26%-55%), and 67% (53%-78%) at the species level for water vole, field vole, and red deer, respectively, the results from 3-6 water replicates would be equivalent to the results from 3-5 latrine surveys and 5-30 weeks of single camera deployments, which provide comparable results to conventional survey methods per unit of survey effort (Sales et al., 2020). In other words, the detection probability of 40% indicates the viability of riverine water eDNA monitoring definite species. Similarity, we set the monitoring effectiveness of 40% to indicate the viability of riverine water eDNA monitoring biodiversity information. In the current case, the effectiveness of monitoring riparian biodiversity information was higher than 40% for all three taxonomic communities at the species level (Table 5). This result suggests that riverine water eDNA is viable for monitoring riparian biodiversity information.

Previous studies have shown that eDNA metabarcoding surveys are relatively cheaper, more efficient, and more accurate than traditional surveys in aquatic systems (Civade, et al., 2016; Valentini, et al., 2016; Lugg, et al., 2018), although this is certainly not true in all circumstances (Beng & Corlett,2020). Considering our work, in which as much as 98.69% bacterial species, 95.71% fungal species, and 92.41% eukaryotic species from 1 km upstream could be detected, we accepted the conclusion that using water eDNA to monitor upstream biodiversity is viable.

Previous studies have shown that the detection distance of eDNA vary from less than 1 km to more than 100 km (Deiner & Altermatt,2014; Stoeckle et al., 2016; Sansom & Sassoubre,2017; Tillotson et al., 2018; Seymour,2019). In our case, the half-life distance of noneffective WBIF for bacteria was approximately 13-19 km and for eukaryotes approximately 4-6 km (Table 6). Thus, we suggested that to efficiently monitor the aquatic and riparian bacterial or eukaryotic communities of the Shaliu River, riverine water eDNA should be sampled on summer and autumn rainy days and the distance between adjacent transects could be set as 13-19 km or 4-6 km, respectively.

## 5 Conclusion

Based on the finding that riverine water eDNA incorporates biodiversity information across terrestrial and aquatic biomes (Deiner et al., 2016; Matsuoka et al., 2019), we propose the idea of using riverine water eDNA to simultaneously monitor aquatic and terrestrial biodiversity. Yes, it is a time- and cost-efficient approach for monitoring watershed biodiversity. To verify the probability of simultaneously monitoring aquatic and riparian biodiversity using riverine water eDNA, we identified the optimal holistic biodiversity monitoring season and weather conditions and studied the taxonomic variation of monitoring effectiveness. Our work showed that (1) summer and autumn were the optimal seasons, and rainy days were the optimal weather condition, as determined by temperature, discharge, flow velocity, and surface runoff; (2) the effectiveness of monitoring bacterial communities was significantly higher than monitoring eukaryotic communities, in which microorganisms left more traces than macroorganisms, prokaryotes left more traces than eukaryotes, eDNA contained inside prokaryotic cells was more resilient against degradation than that contained inside eukaryotic cells; (3) the monitoring effectiveness was significantly higher at the species level than at the OTU level. In the optimal holistic biodiversity monitoring season and weather conditions (a summer rainy day) in a watershed on the Qinghai-Tibet Plateau (the Shaliu river basin), using riverine water eDNA, we detected as much as 87.95% bacterial species, 76.18% fungal species, and 53.52% eukaryotic species from riparian soil, and as much as 98.69% bacterial species, 95.71% fungal species and 92.41% eukaryotic species from 1 km upstream. Our findings support the viability of using riverine water eDNA to simultaneously monitor aquatic and riparian biodiversity.

Considering the half-life distance of noneffective WBIF for bacteria (13-19 km) and eukaryotes (4-6 km), we suggest that to efficiently monitor the aquatic and riparian bacterial or eukaryotic communities of the Shaliu River, riverine water eDNA should be sampled on summer and autumn rainy days and the distance between adjacent transects could be set as 13-19 km or 4-6 km, respectively. In this monitoring program, monitoring the aquatic and riparian biodiversity, we will simply need to sample riverine water eDNA, and riparian soil eDNA samples will no longer be needed, which is time- and cost-efficient. Our findings underscore the potential value of surveying riverine water eDNA as a routine watershed biodiversity monitoring approach to support efficient assessments of aquatic and terrestrial biodiversity. We encourage more studies on the monitoring effectiveness of using riverine water eDNA to monitor aquatic and terrestrial biodiversity for each taxonomic community in other watersheds with different environmental conditions. In future ecological research, biodiversity conservation and ecosystem management, riverine water eDNA may be a general material for routine watershed biodiversity monitoring and assessment.

## Acknowledgements

This work was supported by the Central Public-Interest Scientific Institution Basal Research Fund, Chinese Academy of Fishery Sciences (grant numbers 2019HY-XKQ02, 2020TD08), the Natural Science Foundation of Qinghai (grant numbers 2018-ZJ-908), and the Department of Science and Technology of Qinghai Provence (grant numbers 2018-ZJ-703).

## Data Accessibility Statement

The datasets generated for this study can be found in the China National GeneBank Sequence Archive (CNSA, https://db.cngb.org/cnsa/) of the China National GeneBank database (CNGBdb) under accession number CNP0001046.

## Author Contributions

HY acquired fund, designed and performed research, analyzed data, wrote and edited the paper.

HD acquired fund, designed and supervised research, reviewed and edited the paper.

HQ acquired fund, designed research, reviewed and edited the paper.

LY performed field sampling.

XH performed laboratory experiment.

HZ, JL, JW and CW reviewed and edited the paper.

QZ administrated project.

QW supervised and validated research.

## Supplementary material 1

### Sampling and sequencing

To assess the monitoring effectiveness (approximately indicated by the WBIF transportation effectiveness), we compared the biodiversity information detected in adjacent upstream-to-downstream samples and that detected in adjacent riparian-to-water samples. To identify the seasonal variation of monitoring effectiveness, we sampled three times respectively in Spring, Summer, and Autumn. To identify the taxonomic variation of monitoring effectiveness, we analyzed three taxonomic communities using the metabarcoding of the 16S rRNA gene, ITS gene, and mitochondrial CO1 gene (Collins et al., 2019; Heeger et al., 2019; Giebner et al., 2020). To determine the eDNA of all taxonomic communities, we selected the finest filter membrane (0.2 μm) to filter riverine water samples (Eichmiller, et al., 2016; Li, et al., 2018). Because keeping samples cooled could reduce the rate of eDNA decay and is a convenient and efficient method for conserving eDNA samples (Sales, et al., 2019), we kept our eDNA samples in an ice bath or in a dry ice bath when they were transported and in an ultra-low temperature freezer for storage.

### Field sampling

On April 8 and 9, June 25 and 26, and September 19 and 20, 2019, we collected eDNA samples three times, including 27 soil eDNA samples and 27 water eDNA samples, from 9 transects of the Shaliu River (Fig. 1). A 5-mL soil eDNA sample (actual surface soil) was collected using a 5-mL sterilized centrifuge tube from the riparian site (5 m from the river) of each transect and transported in an ice bath to the laboratory of the Rescue and Rehabilitation Center of Naked Carps of Qinghai Lake at 0°C. Then, the tubes with soil eDNA samples were frozen in a −20°C refrigerator, transported at −20°C (in a dry ice bath), and stored at −80°C (in an ultra-low temperature freezer) until DNA extraction. A 1.5-L surface water sample (actual riverine water) was collected using a 1.5-L sterilized bottle (rinsed three times with sampling water) from the river site of each transect and transported at 0°C (in an ice bath) to the laboratory. Then, water samples were filtered using 0.2-μm membrane filters (JinTeng, Tianjin, PRC) to obtain the eDNA sample in the laboratory (every step follows the general rule of the molecular ecology experiment to control contamination), the membrane filters of each water eDNA sample were placed in a 50-mL sterilized centrifuge tube. Then, the tubes with the membrane filters of water eDNA samples were frozen in a −20°C refrigerator, transported at −20°C (in a dry ice bath), and stored at −80°C (in an ultra-low temperature freezer) until DNA extraction.

In the first sampling period (spring group), during April 8 and 9, the air temperature was −6-8°C, the water temperature was −0.5-0.7°C, the frozen river was starting to thaw, the runoff volume was 1.8-3.9 m^3^/s, the flow velocity was 0.63-1.04 m/s, the soil was still frozen, and both days were cloudy with freezing, heavy winds. On the frozen days of April 8 and 9, the river was clear; on these days, we sampled 18 samples (9 water samples and 9 soil samples) at 9 transects following the downstream-to-upstream direction.

In the second sampling period (summer group), during June 25 and 26, the air temperature was 7-17°C, the water temperature was 4.3-16.4°C, the runoff volume was 29.9-45.5 m^3^/s, and the flow velocity was 0.84-2.03 m/s. On the sunny day of June 25, the river was clear; on this day, we collected 4 samples (2 water samples and 2 soil samples) from the SL1 and SL2 transects (two downstream transects). It began to rain (light rain, less than 10mm) on the night of June 25, and on the rainy day (light rain, less than 10mm) of June 26, the river was turbid (different transect, different turbidity); we collected 14 samples (7 water samples and 7 soil samples) from the last 7 transects following the downstream-to-upstream direction.

In the third sampling period (autumn group) on September 19 and 20, the air temperature was 0-10°C, the water temperature was 0.2-8.8°C, part of the transects started to freeze, the runoff volume was 5.7-12.8 m^3^/s, and the flow velocity was 0.57-0.88 m/s. On the rainy day (light rain, less than 10 mm) of September 19, the downstream river section was turbid, and we collected 4 samples (2 water samples and 2 soil samples) from the SL1 and SL2 transects (two downstream transects). On the cloudy day of September 20, the river was clear, and we collected 14 samples (7 water samples and 7 soil samples) from the last 7 transects following the downstream-to-upstream direction.

### DNA Extraction and Sequence Analysis

As the extraction of eDNA (Hermans et al., 2018; Armbrecht et al., 2020), metabarcoding selection (Collins et al., 2019; Heeger et al., 2019; Giebner et al., 2020), amplification approach and sequencing (Nichols et al., 2018) impact the results of eDNA monitoring, consistent methods should be used for comparisons among samples (Dopheide et al., 2019; Giebner et al., 2020; Nicholson et al., 2020). Commercial eDNA labs can help (Ravindran,2019), in which all approaches (including eDNA extraction, primer synthesis, amplification approach, sequencing and contamination control) could be standard. We send them the samples and select the primers, and they give us the results — a set of sequences, and provide an interactive platform to analyze the sequences. In our work, our samples were processed by Shanghai Majorbio Bio-pharm Technology Co., Ltd (Shanghai, China), and our data were analyzed on the free online platform of Majorbio Cloud Platform (www.majorbio.com). As long DNA fragments show a higher decay rate than short fragments (Jo, et al., 2017; Shogren, et al., 2018), short fragments better reflect community richness than long fragments (Wei, et al., 2018; Jo, et al., 2019). Thus, we restricted the amplified fragment length to 300-500 bp. To identify the taxonomic communities, we selected the primers 338F/806R, ITS1F/ ITS2R, and mlCOIintF/ jgHCO2198R to indicate bacteria, fungi, and eukaryotes, respectively (Collins et al., 2019; Heeger et al., 2019; Giebner et al., 2020).

DNA was extracted from eDNA samples using the FastDNA® SPIN Kit for Soil and the FastPrep® Instrument (MP Biomedicals, Santa Ana,CA) according to the manufacturer’s protocols in the eDNA specific lab. Then, the final DNA concentration and purity were determined using a NanoDrop 2000 UV-vis spectrophotometer (Thermo Scientific, Wilmington, USA), and DNA quality was checked by 1% agarose gel electrophoresis.

Three sets of specific primers with barcode (338F/806R, ITS1F/ITS2R and mlCOIintF/jgHCO2198R) were synthesized. The bacterial 16S rRNA gene was amplified with the primers 338F (5’-ACTCCTACGGGAGGCAGCAG-3’) and 806R (5’-GGACTACHVGGGTWTCTAAT-3’) using a PCR thermocycler system (GeneAmp 9700, ABI, USA) (with blank controls) and the following program: 3 min of denaturation at 95°C; 29 cycles of 30 s at 95°C, 30 s for annealing at 55°C, and 45 s for elongation at 72°C; and a final extension at 72°C for 10 min. The PCR assays were performed in triplicate 20-μL mixtures containing 4 μL of 5× FastPfu Buffer, 2 μL of 2.5 mM dNTPs, 0.8 μL of each primer (5 μM), 0.4 μL of FastPfu Polymerase, 0.2 μL of BSA, and 10 ng of template DNA. The fungal ITS gene was amplified with the primers ITS1F (5’-CTTGGTCATTTAGAGGAAGTAA) and ITS2R (5’-GCTGCGTTCTTCATCGATGC) using a PCR thermocycler system (GeneAmp 9700, ABI, USA) (with blank controls) and the following program: 3 min of denaturation at 95°C; 37 cycles of 30 s at 95°C, 30 s for annealing at 53°C, and 45 s for elongation at 72°C; and a final extension at 72°C for 10 min. The PCR assays were performed in triplicate 20-μL mixtures containing 4 μL of 5× FastPfu Buffer, 2 μL of 2.5 mM dNTPs, 0.8 μL of each primer (5 μM), 0.4 μL of FastPfu Polymerase, 0.2 μL of BSA, and 10 ng of template DNA. The eukaryotic mitochondrial CO1 gene was amplified with the primers mlCOIintF (5’-GGWACWGGWTGAACWGTWTAYCCYCC) and jgHCO2198R (5’-TANACYTCNGGRTGNCCRAARAAYCA) using a PCR thermocycler system (GeneAmp 9700, ABI, USA) (with blank controls) and the following program: 5 min of denaturation at 94°C; 35 cycles of 60 s at 94°C, 120 s for annealing at 47°C, and 60 s for elongation at 72°C; and a final extension at 72°C for 5 min. The PCR assays were performed in triplicate 20-μL mixtures containing 4 μL of 5× FastPfu Buffer, 2 μL of 2.5 mM dNTPs, 0.8 μL of each primer (5 μM), 0.4 μL of FastPfu Polymerase, 0.2 μL of BSA, and 10 ng of template DNA. The PCR products were extracted and further purified using the AxyPrep DNA Gel Extraction Kit (Axygen Biosciences, Union City, CA, USA). The resulting PCR products of the same sample were mixed together and checked by 2% agarose gel electrophoresis.

The PCR product amplicons were quantified using QuantiFluor™ -ST (Promega, U.S.) according to the manufacturer’s protocol. According to the sequencing requirements of each sample, the PCR products were mixed on the basis of the proportions. The standard adaptors provided by Illumina (Illumina, San Diego, USA) were linked to the sequencing regions according to PCR reactions. Adapter dimers were removed using beads. Single-stranded DNA fragments were generated using sodium hydroxide. Sample libraries were pooled in equimolar amounts and subjected to paired-end sequencing on an Illumina MiSeq platform (Illumina, San Diego, USA) according to standard protocols.

Raw fastq files were demultiplexed, quality-filtered by Trimmomatic and merged by FLASH (https://ccb.jhu.edu/software/FLASH/index.shtml). Operational taxonomic units (OTUs) were clustered with a 97% similarity cutoff using UPARSE (http://www.drive5.com/uparse/), and chimeric sequences were identified and removed using UCHIME (http://www.drive5.com/uchime/). The taxonomies of each 16S rRNA, ITS, and CO1 gene sequence were analyzed by the RDP Classifier Bayesian algorithm (http://sourceforge.net/projects/rdp-classifier/) against the Silva132/16S_Bacteria database (http://www.arb-silva.de) using a confidence threshold of 70%, against the Unite8.0/ITS_Fungi database (http://unite.ut.ee/index.php) using a confidence threshold of 70%, and against the nt database (standard database), respectively. The OTU numbers, types, and taxonomic features of the samples were analyzed. Community richness (Chao richness index at OTU level) was examined to determine the variation among the three groups using the Mothur software (https://www.mothur.org/wiki/Download_mothur). The data were analyzed online using the Majorbio Cloud Platform (www.majorbio.com). The raw data have been deposited in the China National GeneBank Sequence Archive (CNSA, https://db.cngb.org/cnsa/) of the China National GeneBank database (CNGBdb) under accession number CNP0001046.

## Supplementary material 2

**Table S1.** Biological information features of the samples of three seasonal groups indicated by the bacterial 16S rRNA gene.

| Seasonal group | Clean sequences | Average lengths of sequences (bp) | Types of OTUs | Biological information features |
| --- | --- | --- | --- | --- |
| Spring group | 1,030,826 | 447.54 | 10,602 | 58 phyla, 141 classes, 424 orders, 782 families, 1,895 genera and 4,537 species |
| Summer group | 968,122 | 416.06 | 13,766 |  |
| Autumn group | 842,317 | 416.32 | 16,500 |  |

The spring group was sampled during April 2019; the summer group was sampled during June 2019; the autumn group was sampled during September 2019.

**Table S2.** Transport capacity, environmental filtration, and transportation effectiveness of watershed biological information flow (WBIF) from the riparian sampling site to the adjacent riverine water sampling site in each sampling transect in three seasons.

|  | Spring group |  |  | Summer group |  |  | Autumn group |  |  |
| --- | --- | --- | --- | --- | --- | --- | --- | --- | --- |
|  | Transport capacity | Environmental filtration | Transportation effectiveness | Transport capacity | Environmental filtration | Transportation effectiveness | Transport capacity | Environmental filtration | Transportation effectiveness |
| S-W_SL6b | 0.099499 | 0.414501 | 0.058256 | 0.504529 | 0.064307 | 0.472084 | 0.566649 | 0.138540 | 0.488145 |
| S-W_SL6 | 0.859828 | 0.383092 | 0.530435 | 0.756467 | 0.048747 | 0.719591 | 0.655770 | 0.153814 | 0.554904 |
| S-W_SL5m | 0.221932 | 0.311564 | 0.152786 | 0.776533 | 0.080882 | 0.713725 | 0.475058 | 0.199631 | 0.380221 |
| S-W_SL5 | 0.226740 | 0.360500 | 0.145000 | 0.655574 | 0.109190 | 0.583992 | 0.597921 | 0.254486 | 0.445759 |
| S-W_SL4b | 0.175403 | 0.401328 | 0.105009 | 0.667238 | 0.107198 | 0.595712 | 0.539223 | 0.121664 | 0.473619 |
| S-W_SL4 | 0.184110 | 0.388603 | 0.112564 | 0.784735 | 0.080409 | 0.721635 | 0.570324 | 0.120697 | 0.501487 |
| S-W_SL3 | 0.205840 | 0.409836 | 0.121480 | 0.649059 | 0.095981 | 0.586761 | 0.610110 | 0.143771 | 0.522394 |
| S-W_SL2 | 0.176972 | 0.414116 | 0.103685 | 0.568211 | 0.222821 | 0.441602 | 0.758528 | 0.269508 | 0.554098 |
| S-W_SL1 | 0.428305 | 0.503238 | 0.212766 | 0.257329 | 0.216037 | 0.201736 | 0.579158 | 0.470276 | 0.306794 |

S-W denotes the WBIF from the soil eDNA sample (riparian zone) to the water eDNA sample (river). SL1, SL2, SL3, SL4, SL4b, SL5m, SL5, SL6, and SL6b denote the sampling transects. S-W\_SL1 denotes the WBIF from the riparian zone to the river at SL1. The spring group, summer group, and autumn group indicate the groups that were sampled in April, June, and September 2019, respectively. Transport capacity indicates the proportion of soil microbes that were transported into the river. Environmental filtration indicates the proportion of soil microbes that could not be kept in rivers. Transportation effectiveness indicates the proportion of soil microbes that could be detected in rivers.

**Table S3.** Accumulative runoff distance and accumulative transportation effectiveness from the first sampling transect in each chain of the upstream-to-downstream watershed biological information flow (WBIF) in three seasons.

| Transport Chain | Cumulative<br>Distance/km | Cumulative Effectiveness |  |  |
| --- | --- | --- | --- | --- |
|  |  | Spring group | Summer group | Autumn group |
| Chain A |  |  |  |  |
| SL6b-SL5 | 9 | 0.521352 | 0.798283 | 0.598885 |
| ##-SL4 | 24.5 | 0.412811 | 0.534830 | 0.471711 |
| ##-SL3 | 48 | 0.368327 | 0.486959 | 0.415639 |
| ##-SL2 | 63 | 0.330961 | 0.383295 | 0.308563 |
| ##-SL1 | 70 | 0.119217 | 0.197425 | 0.257389 |
| Chain B |  |  |  |  |
| SL6-SL5 | 8 | 0.217131 | 0.762412 | 0.605484 |
| ##-SL4 | 23.5 | 0.111554 | 0.590463 | 0.469171 |
| ##-SL3 | 47 | 0.093625 | 0.517251 | 0.408623 |
| ##-SL2 | 62 | 0.067231 | 0.414025 | 0.299731 |
| ##-SL1 | 69 | 0.032371 | 0.192146 | 0.248692 |
| Chain C |  |  |  |  |
| SL5m-SL4 | 8.5 | 0.430074 | 0.765880 | 0.667929 |
| ##-SL3 | 32 | 0.334385 | 0.579820 | 0.540136 |
| ##-SL2 | 47 | 0.228181 | 0.448004 | 0.361037 |
| ##-SL1 | 54 | 0.072555 | 0.196740 | 0.298750 |
| Chain D |  |  |  |  |
| SL4b-SL3 | 23 | 0.521333 | 0.738029 | 0.719376 |
| ##-SL2 | 38 | 0.352000 | 0.555864 | 0.437211 |
| ##-SL1 | 45 | 0.108000 | 0.237335 | 0.335324 |

Chains A, B, C, and D denote the 4 chains of the upstream-to-downstream WBIF. In Chain A, SL6b-SL5 denotes the WBIF from sampling transects SL6b to SL5; ##-SL4 denotes the WBIF from sampling transects SL6b to SL5, and then to SL4; ##-SL3 denotes the WBIF from sampling transects SL6b to SL5, SL4, and then to SL3; ##-SL2 denotes the WBIF from sampling transects SL6b to SL5, SL4, SL3, and then to SL2; ##-SL1 denotes the WBIF from sampling transects SL6b to SL5, SL4, SL3, SL2, and then to SL1 (estuary). The spring group, summer group, and autumn group indicate the groups that were sampled in April, June, and September 2019, respectively.

**Table S4.** Biological information features of the samples of three taxonomic groups sampled on summer rainy days.

| Taxonomic group (gene) | Clean sequences | Average lengths of sequences (bp) | Types of OTUs | Biological information features |
| --- | --- | --- | --- | --- |
| 16S rRNA gene | 968,122 | 416.06 | 13,766 | 51 phyla, 128 classes, 380 orders, 687 families, 1,512 genera, 3,532 species |
| ITS gene | 1,287,852 | 237.87 | 7,098 | 15 phyla, 53 classes, 137 orders, 301 families, 644 genera, 1,032 species |
| CO1 gene | 1,283,110 | 317.02 | 17,316 | 43 phyla, 140 classes, 492 orders, 1765 families, 4113 genera, 6836 species |

The 16S rRNA gene, ITS gene, and CO1 gene indicate the taxonomic groups detected by the 16S rRNA gene, ITS gene, and CO1 gene, respectively.

**Table S5.**
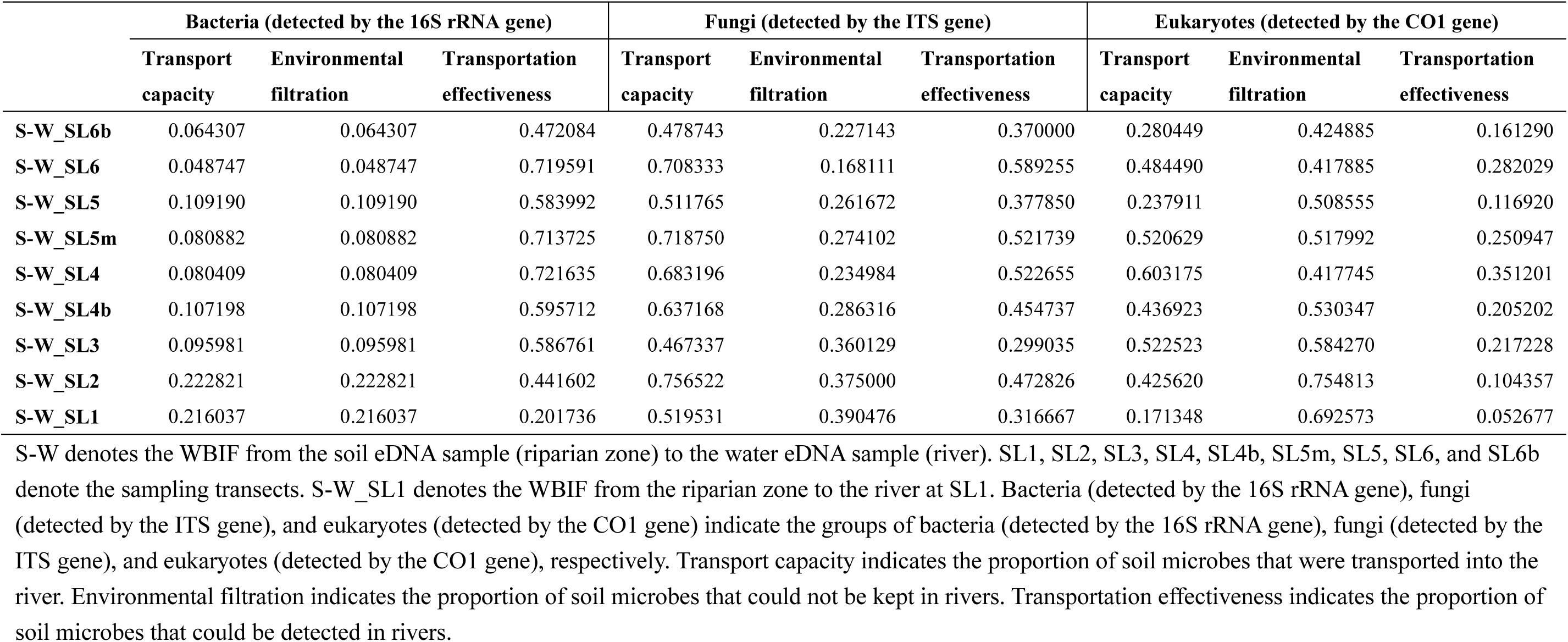
Transport capacity, environmental filtration, and transportation effectiveness of watershed biological information flow (WBIF) from the riparian sampling site to adjacent riverine water sampling site in each sampling transect on summer rainy days, estimated at the OTU level.

|  | Bacteria (detected by the 16S rRNA gene) |  |  | Fungi (detected by the ITS gene) |  |  | Eukaryotes (detected by the CO1 gene) |  |  |
| --- | --- | --- | --- | --- | --- | --- | --- | --- | --- |
|  | Transport capacity | Environmental filtration | Transportation effectiveness | Transport capacity | Environmental filtration | Transportation effectiveness | Transport capacity | Environmental filtration | Transportation effectiveness |
| **S-W_SL6b** | 0.064307 | 0.064307 | 0.472084 | 0.478743 | 0.227143 | 0.370000 | 0.280449 | 0.424885 | 0.161290 |
| **S-W_SL6** | 0.048747 | 0.048747 | 0.719591 | 0.708333 | 0.168111 | 0.589255 | 0.484490 | 0.417885 | 0.282029 |
| **S-W_SL5** | 0.109190 | 0.109190 | 0.583992 | 0.511765 | 0.261672 | 0.377850 | 0.237911 | 0.508555 | 0.116920 |
| **S-W_SL5m** | 0.080882 | 0.080882 | 0.713725 | 0.718750 | 0.274102 | 0.521739 | 0.520629 | 0.517992 | 0.250947 |
| **S-W_SL4** | 0.080409 | 0.080409 | 0.721635 | 0.683196 | 0.234984 | 0.522655 | 0.603175 | 0.417745 | 0.351201 |
| **S-W_SL4b** | 0.107198 | 0.107198 | 0.595712 | 0.637168 | 0.286316 | 0.454737 | 0.436923 | 0.530347 | 0.205202 |
| **S-W_SL3** | 0.095981 | 0.095981 | 0.586761 | 0.467337 | 0.360129 | 0.299035 | 0.522523 | 0.584270 | 0.217228 |
| **S-W_SL2** | 0.222821 | 0.222821 | 0.441602 | 0.756522 | 0.375000 | 0.472826 | 0.425620 | 0.754813 | 0.104357 |
| **S-W_SL1** | 0.216037 | 0.216037 | 0.201736 | 0.519531 | 0.390476 | 0.316667 | 0.171348 | 0.692573 | 0.052677 |

S-W denotes the WBIF from the soil eDNA sample (riparian zone) to the water eDNA sample (river). SL1, SL2, SL3, SL4, SL4b, SL5m, SL5, SL6, and SL6b denote the sampling transects. S-W\_SL1 denotes the WBIF from the riparian zone to the river at SL1. Bacteria (detected by the 16S rRNA gene), fungi (detected by the ITS gene), and eukaryotes (detected by the CO1 gene) indicate the groups of bacteria (detected by the 16S rRNA gene), fungi (detected by the ITS gene), and eukaryotes (detected by the CO1 gene), respectively. Transport capacity indicates the proportion of soil microbes that were transported into the river. Environmental filtration indicates the proportion of soil microbes that could not be kept in rivers. Transportation effectiveness indicates the proportion of soil microbes that could be detected in rivers.

**Table S6.** Transport capacity, environmental filtration, and transportation effectiveness of watershed biological information flow (WBIF) from the riparian sampling site to adjacent riverine water sampling site in each sampling transect on summer rainy days, estimated at the species level.

|  | Bacteria (detected by the 16S rRNA gene) |  |  | Fungi (detected by the ITS gene) |  |  | Eukaryotes (detected by the CO1 gene) |  |  |
| --- | --- | --- | --- | --- | --- | --- | --- | --- | --- |
|  | Transport capacity | Environmental filtration | Transportation effectiveness | Transport capacity | Environmental filtration | Transportation effectiveness | Transport capacity | Environmental filtration | Transportation effectiveness |
| **S-W_SL6b** | 0.683468 | 0.021696 | 0.668639 | 0.625000 | 0.111111 | 0.555556 | 0.437819 | 0.245501 | 0.330334 |
| **S-W_SL6** | 0.874088 | 0.012613 | 0.863063 | 0.827907 | 0.092827 | 0.751055 | 0.600000 | 0.286492 | 0.428105 |
| **S-W_SL5** | 0.844639 | 0.033827 | 0.816068 | 0.591503 | 0.133144 | 0.512748 | 0.504570 | 0.313676 | 0.346299 |
| **S-W_SL5m** | 0.893401 | 0.026680 | 0.869565 | 0.850467 | 0.085470 | 0.777778 | 0.664207 | 0.306905 | 0.460358 |
| **S-W_SL4** | 0.880989 | 0.029257 | 0.855214 | 0.833948 | 0.117264 | 0.736156 | 0.700000 | 0.245283 | 0.528302 |
| **S-W_SL4b** | 0.814159 | 0.032120 | 0.788009 | 0.774359 | 0.129464 | 0.674107 | 0.619165 | 0.257299 | 0.459854 |
| **S-W_SL3** | 0.818637 | 0.032946 | 0.791667 | 0.669643 | 0.125000 | 0.585938 | 0.704082 | 0.313084 | 0.483645 |
| **S-W_SL2** | 0.760110 | 0.074043 | 0.703830 | 0.828244 | 0.176101 | 0.682390 | 0.627685 | 0.433784 | 0.355405 |
| **S-W_SL1** | 0.442045 | 0.081420 | 0.406054 | 0.594118 | 0.162562 | 0.497537 | 0.271255 | 0.433486 | 0.153670 |

S-W denotes the WBIF from the soil eDNA sample (riparian zone) to the water eDNA sample (river). SL1, SL2, SL3, SL4, SL4b, SL5m, SL5, SL6, and SL6b denote the sampling transects. S-W\_SL1 denotes the WBIF from the riparian zone to the river at SL1. Bacteria (detected by the 16S rRNA gene), fungi (detected by the ITS gene), and eukaryotes (detected by the CO1 gene) indicate the groups of bacteria (detected by the 16S rRNA gene), fungi (detected by the ITS gene), and eukaryotes (detected by the CO1 gene), respectively. Transport capacity indicates the proportion of soil microbes that were transported into the river. Environmental filtration indicates the proportion of soil microbes that could not be kept in rivers. Transportation effectiveness indicates the proportion of soil microbes that could be detected in rivers.

**Table S7.**
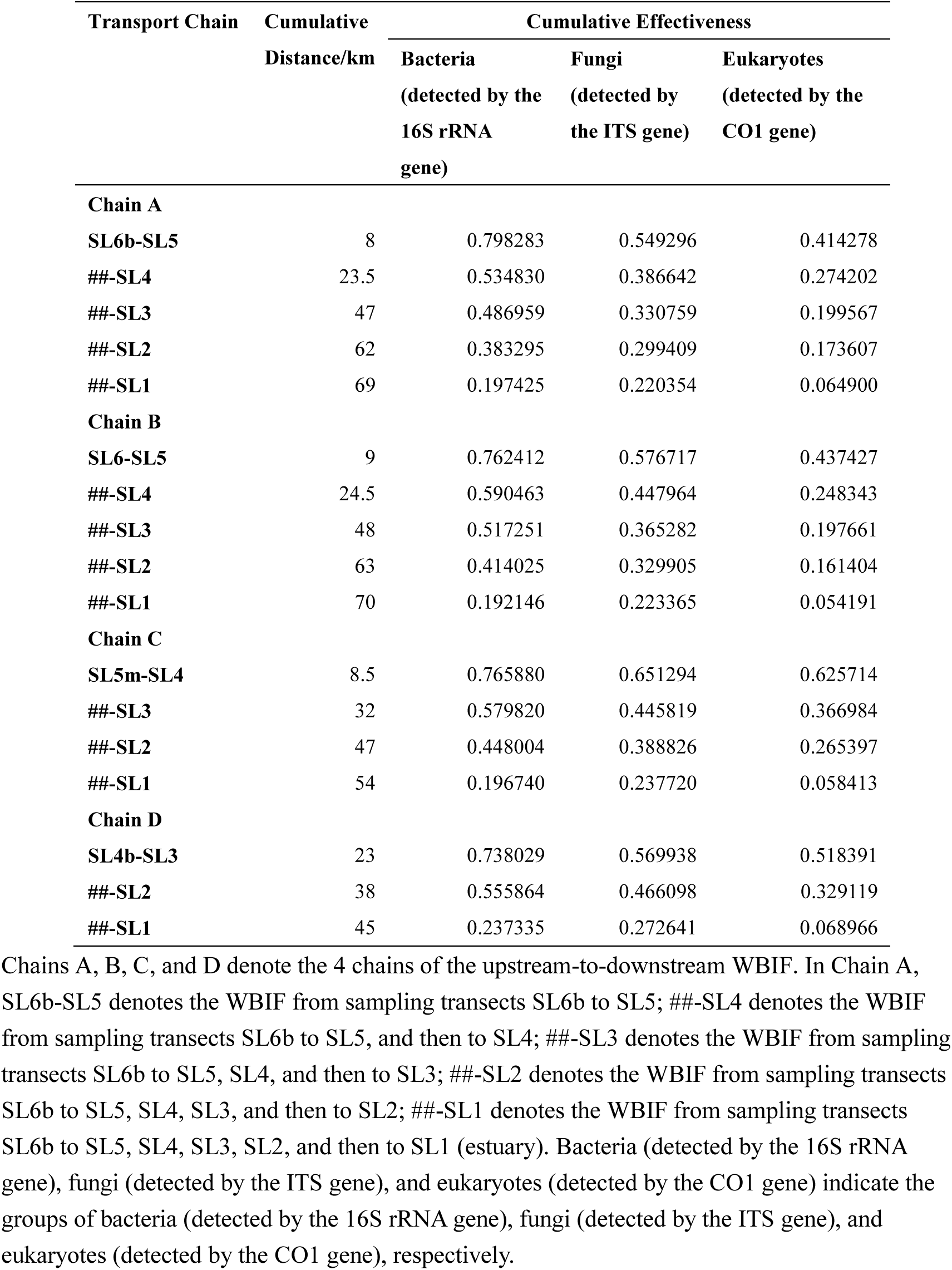
The accumulative runoff distance and accumulative transportation effectiveness from the first sampling transect in each chain of the upstream-to-downstream watershed biological information flow (WBIF) on summer rainy days, estimated at the OTU level.

**Table S8.** The accumulative runoff distance and accumulative transportation effectiveness from the first sampling transect in each chain of the upstream-to-downstream watershed biological information flow (WBIF) on summer rainy days, estimated at the species level.

| Transport Chain | Cumulative<br>Distance/km | Cumulative Effectiveness |  |  |
| --- | --- | --- | --- | --- |
|  |  | Bacteria | Fungi | Eukaryotes |
|  |  | (detected by the<br>16S rRNA gene) | (detected by<br>the ITS gene) | (detected by the<br>CO1 gene) |
| Chain A |  |  |  |  |
| SL6b-SL5 | 8 | 0.897101 | 0.757895 | 0.515347 |
| ##-SL4 | 23.5 | 0.759420 | 0.701754 | 0.384491 |
| ##-SL3 | 47 | 0.711594 | 0.677193 | 0.312601 |
| ##-SL2 | 62 | 0.610870 | 0.659649 | 0.280291 |
| ##-SL1 | 69 | 0.377536 | 0.557895 | 0.137318 |
| Chain B |  |  |  |  |
| SL6-SL5 | 9 | 0.862913 | 0.634091 | 0.501928 |
| ##-SL4 | 24.5 | 0.775398 | 0.590909 | 0.322865 |
| ##-SL3 | 48 | 0.712974 | 0.534091 | 0.271074 |
| ##-SL2 | 63 | 0.616891 | 0.522727 | 0.229201 |
| ##-SL1 | 70 | 0.355569 | 0.422727 | 0.099725 |
| Chain C |  |  |  |  |
| SL5m-SL4 | 8.5 | 0.868212 | 0.787819 | 0.670061 |
| ##-SL3 | 32 | 0.753508 | 0.595285 | 0.454684 |
| ##-SL2 | 47 | 0.640635 | 0.563851 | 0.341650 |
| ##-SL1 | 54 | 0.356925 | 0.408644 | 0.112016 |
| Chain D |  |  |  |  |
| SL4b-SL3 | 23 | 0.848338 | 0.688679 | 0.601319 |
| ##-SL2 | 38 | 0.715374 | 0.646226 | 0.414868 |
| ##-SL1 | 45 | 0.393352 | 0.457547 | 0.126499 |

Chains A, B, C, and D denote the 4 chains of the upstream-to-downstream WBIF. In Chain A, SL6b-SL5 denotes the WBIF from sampling transects SL6b to SL5; ##-SL4 denotes the WBIF from sampling transects SL6b to SL5, and then to SL4; ##-SL3 denotes the WBIF from sampling transects SL6b to SL5, SL4, and then to SL3; ##-SL2 denotes the WBIF from sampling transects SL6b to SL5, SL4, SL3, and then to SL2; ##-SL1 denotes the WBIF from sampling transects SL6b to SL5, SL4, SL3, SL2, and then to SL1 (estuary). Bacteria (detected by the 16S rRNA gene), fungi (detected by the ITS gene), and eukaryotes (detected by the CO1 gene) indicate the groups of bacteria (detected by the 16S rRNA gene), fungi (detected by the ITS gene), and eukaryotes (detected by the CO1 gene), respectively.

## Footnotes

1 http://www.gangcha.gov.cn/html/2125/item.html

## References

Altermatt F, Little CJ, Mächler E, et al. (2020) Uncovering the complete biodiversity structure in spatial networks: the example of riverine systems. Oikos. https://doi.org/10.1111/oik.06806

Anderson CB (2018) Biodiversity monitoring, earth observations and the ecology of scale. Ecology Letters 21, 1572–1585. https://doi.org/10.1111/ele.13106

Armbrecht, L. & S. Herrando Pérez, et al. (2020). An optimized method for the extraction of ancient eukaryote DNA from marine sediments. Molecular Ecology Resources 20, 906–919. https://doi.org/10.1111/1755-0998.13162

Bálint, M. & M. Pfenninger, et al. (2018). Environmental DNA Time Series in Ecology. Trends in Ecology & Evolution 33, 945–957. https://doi.org/10.1016/j.tree.2018.09.003

Barnes, M. A. & C. R. Turner, et al. (2014). Environmental Conditions Influence eDNA Persistence in Aquatic Systems. Environmental Science & Technology 48, 1819–1827. https://doi.org/10.1021/es404734p

Barnes MA, Turner CR (2016) The ecology of environmental DNA and implications for conservation genetics. Conservation Genetics 17, 1–17. https://doi.org/10.1007/s10592-015-0775-4

Bass, D. & G. D. Stentiford, et al. (2015). Diverse Applications of Environmental DNA Methods in Parasitology. Trends in Parasitology 31, 499–513. https://doi.org/10.1016/j.pt.2015.06.013

Beng, K. C. & R. T. Corlett (2020). Applications of environmental DNA (eDNA) in ecology and conservation: opportunities, challenges and prospects. Biodiversity and Conservation 29, 2089–2121. https://doi.org/10.1007/s10531-020-01980-0

Cardinale BJ, Duffy JE, Gonzalez A, et al. (2012) Biodiversity loss and its impact on humanity. Nature 486, 59–67. https://doi.org/10.1038/nature11148

Carraro L, Hartikainen H, Jokela J, Bertuzzo E, Rinaldo A (2018) Estimating species distribution and abundance in river networks using environmental DNA. Proceedings of the National Academy of Sciences of the United States of America 115, 11724–11729. https://doi.org/10.1073/pnas.1813843115

Carraro, L. & E. Mächler, et al. (2020). Environmental DNA allows upscaling spatial patterns of biodiversity in freshwater ecosystems. Nature Communications 11, 3585. https://doi.org/10.1038/s41467-020-17337-8

Civade, R. & T. Dejean, et al. (2016). Spatial Representativeness of Environmental DNA Metabarcoding Signal for Fish Biodiversity Assessment in a Natural Freshwater System. PLoS One 11, e0157366–19. https://doi.org/10.1371/journal.pone.0157366

Collins, R. A. & J. Bakker, et al. (2019). Non-specific amplification compromises environmental DNA metabarcoding with COI. Methods in Ecology and Evolution 10, 1985–2001. https://doi.org/10.1111/2041-210X.13276

Cristescu ME, Hebert PDN (2018) Uses and Misuses of Environmental DNA in Biodiversity Science and Conservation. Annual Review of Ecology, Evolution, and Systematics 49, 209–230. https://doi.org/10.1146/annurev-ecolsys-110617-062306

Deiner K, Altermatt F (2014) Transport Distance of Invertebrate Environmental DNA in a Natural River. PLoS ONE 9, e88786. https://doi.org/10.1371/journal.pone.0088786

Deiner K, Fronhofer EA, Mächler E, Walser J, Altermatt F (2016) Environmental DNA reveals that rivers are conveyer belts of biodiversity information. Nature Communications 7, 12544. https://doi.org/10.1038/ncomms12544

Dixon, K. M. & G. J. Cary, et al. (2019). Features associated with effective biodiversity monitoring and evaluation. Biological Conservation 238, 108221. https://doi.org/10.1016/j.biocon.2019.108221

Djurhuus, A. & C. J. Closek, et al. (2020). Environmental DNA reveals seasonal shifts and potential interactions in a marine community. Nature Communications 11, 254. https://doi.org/10.1038/s41467-019-14105-1

Dopheide, A. & D. Xie, et al. (2019). Impacts of DNA extraction and PCR on DNA metabarcoding estimates of soil biodiversity. Methods in Ecology and Evolution 10, 120–133. https://doi.org/10.1111/2041-210X.13086

Eichmiller, J. J. & S. E. Best, et al. (2016). Effects of temperature and trophic state on degradation of environmental DNA in lake water. Environmental Science & Technology 50, 1859–1867. https://doi.org/10.1021/acs.est.5b05672

Fremier, A. K. & K. M. Strickler, et al. (2019). Stream transport and retention of environmental DNA pulse releases in relation to hydrogeomorphic scaling factors. Environmental Science & Technology 53, 6640–6649. https://doi.org/10.1021/acs.est.8b06829

Giebner, H. & K. Langen, et al. (2020). Comparing diversity levels in environmental samples: DNA sequence capture and metabarcoding approaches using 18S and COI genes. Molecular Ecology Resources 20, 1333–1345. https://doi.org/10.1111/1755-0998.13201

Gogarten, J. F. & C. Hoffmann, et al. (2019). Fly-derived DNA and camera traps are complementary tools for assessing mammalian biodiversity. Environmental DNA 2, 63–76. https://doi.org/10.1002/edn3.46

Gusareva, E. S. & E. Acerbi, et al. (2019). Microbial communities in the tropical air ecosystem follow a precise diel cycle. Proceedings of the National Academy of Sciences 116, 23299–23308. https://doi.org/10.1073/pnas.1908493116

Harper, L. R. & L. Lawson Handley, et al. (2019). Environmental DNA (eDNA) metabarcoding of pond water as a tool to survey conservation and management priority mammals. Biological Conservation 238, 108225. https://doi.org/10.1016/j.biocon.2019.108225

Heeger, F. & C. Wurzbacher, et al. (2019). Combining the 5.8S and ITS2 to improve classification of fungi. Methods in Ecology and Evolution 10, 1702–1711. https://doi.org/10.1111/2041-210X.13266

Hermans, S. M. & H. L. Buckley, et al. (2018). Optimal extraction methods for the simultaneous analysis of DNA from diverse organisms and sample types. Molecular Ecology Resources 18, 557–569. https://doi.org/10.1111/1755-0998.12762

Hooper DU, Adair EC, Cardinale BJ, et al. (2012) A global synthesis reveals biodiversity loss as a major driver of ecosystem change. Nature 486, 105–108. https://doi.org/10.1038/nature11118

Jerde, C. L. & B. P. Olds, et al. (2016). Influence of stream bottom substrate on retention and transport of vertebrate environmental DNA. Environmental Science & Technology 50, 8770–8779. https://doi.org/10.1021/acs.est.6b01761

Jetz W, Cavender-Bares J, Pavlick R, et al. (2016) Monitoring plant functional diversity from space. Nature Plants 2, 16024. https://doi.org/10.1038/nplants.2016.24

Jo, T. & H. Murakami, et al. (2017). Rapid degradation of longer DNA fragments enables the improved estimation of distribution and biomass using environmentalDNA. Molecular Ecology Resources 17, e25–e33. https://doi.org/10.1111/1755-0998.12685

Jo, T. & M. Arimoto, et al. (2019). Particle Size Distribution of Environmental DNA from the Nuclei of Marine Fish. Environmental Science & Technology 53, 9947–9956. https://doi.org/10.1021/acs.est.9b02833

Johnson, M. D. & R. D. Cox, et al. (2019). Analyzing airborne environmental DNA: A comparison of extraction methods, primer type, and trap type on the ability to detect airborne eDNA from terrestrial plant communities. Environmental DNA 1, 176–185. https://doi.org/10.1002/edn3.19

Koranda, M. & C. Kaiser, et al. (2013). Seasonal variation in functional properties of microbial communities in beech forest soil. Soil Biology and Biochemistry 60, 95–104. https://doi.org/10.1016/j.soilbio.2013.01.025

Li, J. & L. Lawson Handley, et al. (2018). The effect of filtration method on the efficiency of environmental DNA capture and quantification via metabarcoding. Molecular Ecology Resources 18, 1102–1114. https://doi.org/10.1111/1755-0998.12899

Lin, Q. & J. De Vrieze, et al. (2016). Temperature affects microbial abundance, activity and interactions in anaerobic digestion. Bioresource Technology 209, 228–236. https://doi.org/10.1016/j.biortech.2016.02.132

Lugg WH, Griffiths J, van Rooyen AR, Weeks AR, Tingley R (2018) Optimal survey designs for environmental DNA sampling. Methods in Ecology and Evolution 9, 1049–1059. https://doi.org/10.1111/2041-210X.12951

Luo Y, Xu L, Rysz M, et al. (2011) Occurrence and Transport of Tetracycline, Sulfonamide, Quinolone, and Macrolide Antibiotics in the Haihe River Basin, China. Environmental Science & Technology 45, 1827–1833. https://doi.org/10.1021/es104009s

Matsuoka S, Sugiyama Y, Sato H, et al. (2019) Spatial structure of fungal DNA assemblages revealed with eDNA metabarcoding in a forest river network in western Japan. Metabarcoding and Metagenomics 3: e36335. https://doi.org/10.3897/mbmg.3.36335

Nichols, R. V. & C. Vollmers, et al. (2018). Minimizing polymerase biases in metabarcoding. Molecular Ecology Resources 18, 927–939. https://doi.org/10.1111/1755-0998.12895

Nicholson, A. & D. McIsaac, et al. (2020). An analysis of metadata reporting in freshwater environmental DNA research calls for the development of best practice guidelines. Environmental DNA 2, 343–349. https://doi.org/10.1002/edn3.81

Nukazawa, K. & Y. Hamasuna, et al. (2018). Simulating the advection and degradation of the environmental DNA of common carp along a river. Environmental Science & Technology 52, 10562–10570. https://doi.org/10.1021/acs.est.8b02293

Pawlowski, J. & L. Apothéloz-Perret-Gentil, et al. (2020). Environmental DNA: What’s behind the term? Clarifying the terminology and recommendations for its future use in biomonitoring. Molecular Ecology 29, 4258–4264. https://doi.org/10.1111/mec.15643

Pont D, Rocle M, Valentini A, et al. (2018) Environmental DNA reveals quantitative patterns of fish biodiversity in large rivers despite its downstream transportation. Scientific Reports 8, 10313–10361. https://doi.org/10.1038/s41598-018-28424-8

Ravindran, S. (2019). Turning discarded DNA into ecology gold. Nature 570, 543–545. https://doi.org/10.1038/d41586-019-01987-w

Rodgers, T. W. & K. E. Mock (2015). Drinking water as a source of environmental DNA for the detection of terrestrial wildlife species. Conservation Genetics Resources 7, 693–696. https://doi.org/10.1007/s12686-015-0478-7

Sales, N. G. & O. S. Wangensteen, et al. (2019). Influence of preservation methods, sample medium and sampling time on eDNA recovery in a neotropical river. Environmental DNA 1, 119–130. https://doi.org/10.1002/edn3.14

Sales NG, McKenzie MB, Drake J, et al. (2020) Fishing for mammals: Landscape-level monitoring of terrestrial and semi-aquatic communities using eDNA from riverine systems. Journal of Applied Ecology 57, 707–716. https://doi.org/10.1111/1365-2664.13592

Sansom BJ, Sassoubre LM (2017) Environmental DNA (eDNA) Shedding and Decay Rates to Model Freshwater Mussel eDNA Transport in a River. Environmental Science & Technology 51, 14244–14253. https://doi.org/10.1021/acs.est.7b05199

Seeber, P. A. & G. K. McEwen, et al. (2019). Terrestrial mammal surveillance using hybridization capture of environmental DNA from African waterholes. Molecular Ecology Resources 19, 1486–1496. https://doi.org/10.1111/1755-0998.13069

Seymour M (2019) Rapid progression and future of environmental DNA research. Communications Biology 2, 80. https://doi.org/10.1038/s42003-019-0330-9

Seymour, M. & I. Durance, et al. (2018). Acidity promotes degradation of multi-species environmental DNA in lotic mesocosms. Communications Biology 1, 4. https://doi.org/10.1038/s42003-017-0005-3

Shogren AJ, Tank JL, Andruszkiewicz E, et al. (2017) Controls on eDNA movement in streams: Transport, Retention, and Resuspension. Scientific Reports 7, 5065. https://doi.org/10.1038/s41598-017-05223-1

Shogren, A. J. & J. L. Tank, et al. (2018). Water flow and biofilm cover influence environmental DNA detection in recirculating streams. Environmental Science & Technology 52, 8530–8537. https://doi.org/10.1021/acs.est.8b01822

Shogren, A. J. & J. L. Tank, et al. (2019). Riverine distribution of mussel environmental DNA reflects a balance among density, transport, and removal processes. Freshwater Biology 64, 1467–1479. https://doi.org/10.1111/fwb.13319

Thomsen PF, Willerslev E (2015) Environmental DNA – An emerging tool in conservation for monitoring past and present biodiversity. Biological Conservation 183, 4–18. https://doi.org/10.1016/j.biocon.2014.11.019

Tillotson MD, Kelly RP, Duda JJ, et al. (2018) Concentrations of environmental DNA (eDNA) reflect spawning salmon abundance at fine spatial and temporal scales. Biological Conservation 220, 1–11. https://doi.org/10.1016/j.biocon.2018.01.030

Ushio, M. & H. Fukuda, et al. (2017). Environmental DNA enables detection of terrestrial mammals from forest pond water. Molecular Ecology Resources 17, e63–e75. https://doi.org/10.1111/1755-0998.12690

Valentini A, Taberlet P, Miaud C, et al. (2016) Next-generation monitoring of aquatic biodiversity using environmental DNA metabarcoding. Molecular Ecology 25, 929–942. https://doi.org/10.1111/mec.13428

van Bochove K, Bakker FT, Beentjes KK, et al. (2020) Organic matter reduces the amount of detectable environmental DNA in freshwater. Ecology and Evolution 10, 3647–3654. https://doi.org/10.1002/ece3.6123

Wacker, S. & F. Fossøy, et al. (2019). Downstream transport and seasonal variation in freshwater pearl mussel (Margaritifera margaritifera) eDNA concentration. Environmental DNA 1, 64–73. https://doi.org/10.1002/edn3.10

Wang S, Fu B, Piao S, et al. (2016) Reduced sediment transport in the Yellow River due to anthropogenic changes. Nature Geoscience 9, 38–41. https://doi.org/10.1038/ngeo2602

Wei, N. & F. Nakajima, et al. (2018). A Microcosm Study of Surface Sediment Environmental DNA: Decay Observation, Abundance Estimation, and Fragment Length Comparison. Environmental Science & Technology 52, 12428–12435. https://doi.org/10.1021/acs.est.8b04956

Wilcox, T. M. & K. E. Zarn, et al. (2018). Capture enrichment of aquatic environmental DNA: A first proof of concept. Molecular Ecology Resources 18, 1392–1401. https://doi.org/10.1111/1755-0998.12928

Wilpiszeski, R. L. & J. A. Aufrecht, et al. (2019). Soil aggregate microbial communities: towards understanding microbiome interactions at biologically relevant scales. Applied and Environmental Microbiology 85, e00324–19. https://doi.org/10.1128/aem.00324-19

Xu, B. & J. Wang, et al. (2018). Seasonal and interannual dynamics of soil microbial biomass and available nitrogen in an alpine meadow in the eastern part of Qinghai–Tibet Plateau, China. Biogeosciences 15, 567–579. https://doi.org/10.5194/bg-15-567-2018

